# *De novo* design of hundreds of functional GPCR-targeting antibodies enabled by scaling test-time compute

**DOI:** 10.1101/2025.05.28.656709

**Authors:** Nabla Bio, Surojit Biswas

## Abstract

We present significant advances in *de novo* antibody design against G protein-coupled receptors (GPCRs) enabled by scaling the test-time compute used by our generative protein design system, JAM. We *de novo* design hundreds of VHH (single domain) antibodies against CXCR4 and CXCR7, with top designs showing picomolar to low-nanomolar affinities, high selectivity, and favorable early-stage developability profiles, matching or outperforming clinical-stage molecules in these dimensions. Further, high affinity designs potently modulate receptor function, with most acting as antagonists (inhibitors) and, strikingly, a subset functioning as agonists (activators) of CXCR7 — the first antibody agonists reported for this receptor, and the first computationally designed antibody GPCR agonists of any kind. Using a single experimentally validated agonist to further prompt JAM, we generate over 300 additional diverse agonists with superior properties, including a design with agonism EC50 rivaling that of CXCR7’s natural ligand, SDF1α. These results show that increasing the “reasoning” capacity of biomolecular generative models by scaling test-time computation will enable them to solve increasingly difficult problems in drug design.

## Introduction

Recent advances in large language models (LLMs) have demonstrated that increasing computational resources at application or inference time — a concept known as "test-time scaling" — can dramatically improve model performance without additional training^1,2^. By enabling models to "think longer" through techniques such as iterative reasoning and solution refinement, test-time scaling has allowed LLMs to solve increasingly complex problems by exploring solution spaces more thoroughly^3,4^. This has established a new frontier of performance where models can tackle challenges previously thought intractable — from solving complex mathematical proofs to performing multi-step reasoning in scientific domains — without requiring retraining or larger architectures^5^.

While significant research has focused on scaling laws for pretraining biomolecular generative models — examining how model size, training data, and parameter count affect performance^6–9^— the potential of test-time scaling remains relatively unexplored in computational protein design.

G protein-coupled receptors (GPCRs) represent one of the most therapeutically important protein families, mediating diverse physiological processes and serving as targets for over one-third of FDA-approved drugs^10,11^. Despite their clinical significance, most GPCR-targeting therapeutics are small molecules or peptides that often lack selectivity, leading to off-target effects^12^. Antibodies offer compelling advantages including higher specificity, longer half-lives, and peripheral restriction, and yet have been largely restricted from the GPCR therapeutic landscape due to challenges in integrating these complex membrane proteins into traditional antibody discovery workflows^13,14,15^. Beyond merely binding, modulating GPCR signaling offers control over key biological pathways, making the generation of functional antibodies against GPCRs one of the most compelling opportunities, but persistent challenges in therapeutic discovery today^13^.

Recent work has shown promising results for computational design of mini-proteins against GPCRs, suggesting deep learning based design methodologies can address these difficult targets despite their conformational flexibility and membrane-bound nature^16^. However, similarly successful approaches for antibodies have remained elusive. In our previous work, we introduced JAM (Joint Atomic Modeling), a generative *de novo* antibody design system capable of creating antibodies against diverse targets, including the first fully computationally designed antibody against a GPCR^17^. However, this singular design showed only modest affinity, poor developability, and lacked functional activity, highlighting a gap between feasibility and therapeutic relevance.

We hypothesized that applying test-time scaling to JAM could bridge this gap, just as test-time scaling has empowered language models to solve increasingly difficult reasoning tasks. By allowing JAM more computational resources during inference to explore and refine potential solutions before testing them experimentally, we aimed to generate *de novo* designs with superior binding properties and functional capabilities for GPCRs.

In this work, we apply test-time scaling to *de novo* antibody design through an approach we term "introspection," where JAM progressively refines antibody designs through multiple computational rounds prior to wet-lab testing. This approach yields 20-70x improvements in binding success rates for VHH antibodies targeting SARS-CoV-2 RBD and the GPCRs CXCR4 and CXCR7. Our top designs exhibit picomolar to low-nanomolar affinities with high selectivity, favorable developability profiles, and therapeutically relevant functional modulation. While most designs inhibit GPCR activity — with CXCR4-targeting designs outperforming a clinical-stage benchmark — strikingly, a subset activates CXCR7, representing the first antibody agonists reported for this receptor. Using a single experimentally validated agonist to re-prompt JAM — an approach we refer to as *experiment-guided steering* — we efficiently generate 300+ diverse improved agonists, some with potency rivaling CXCR7’s natural ligand. These findings highlight test-time scaling’s potential to increase the scope of drug design problems solvable with generative protein design.

## Results

### Test-time scaling improves *de novo* VHH affinities and success rates by >10-fold

In recent work, we introduced JAM, a general-purpose generative protein design system whose input and output spaces are protein complexes^17,18^. We define a protein complex as the sequence and all-atom structure of either a single protein chain or a collection of physically interacting protein chains. Given partially specified sequence and structure information for a protein complex, JAM generatively "fills in" unknown parts (Fig. 1a). For example, in this work, we use JAM to generate an antibody’s sequence and structure given target sequence, structure, and epitope. We also use JAM to generate soluble and stable protein scaffolds that preserve and present the extracellular epitopes of GPCRs, which together function as antibody screening reagents.

**Figure 1.**
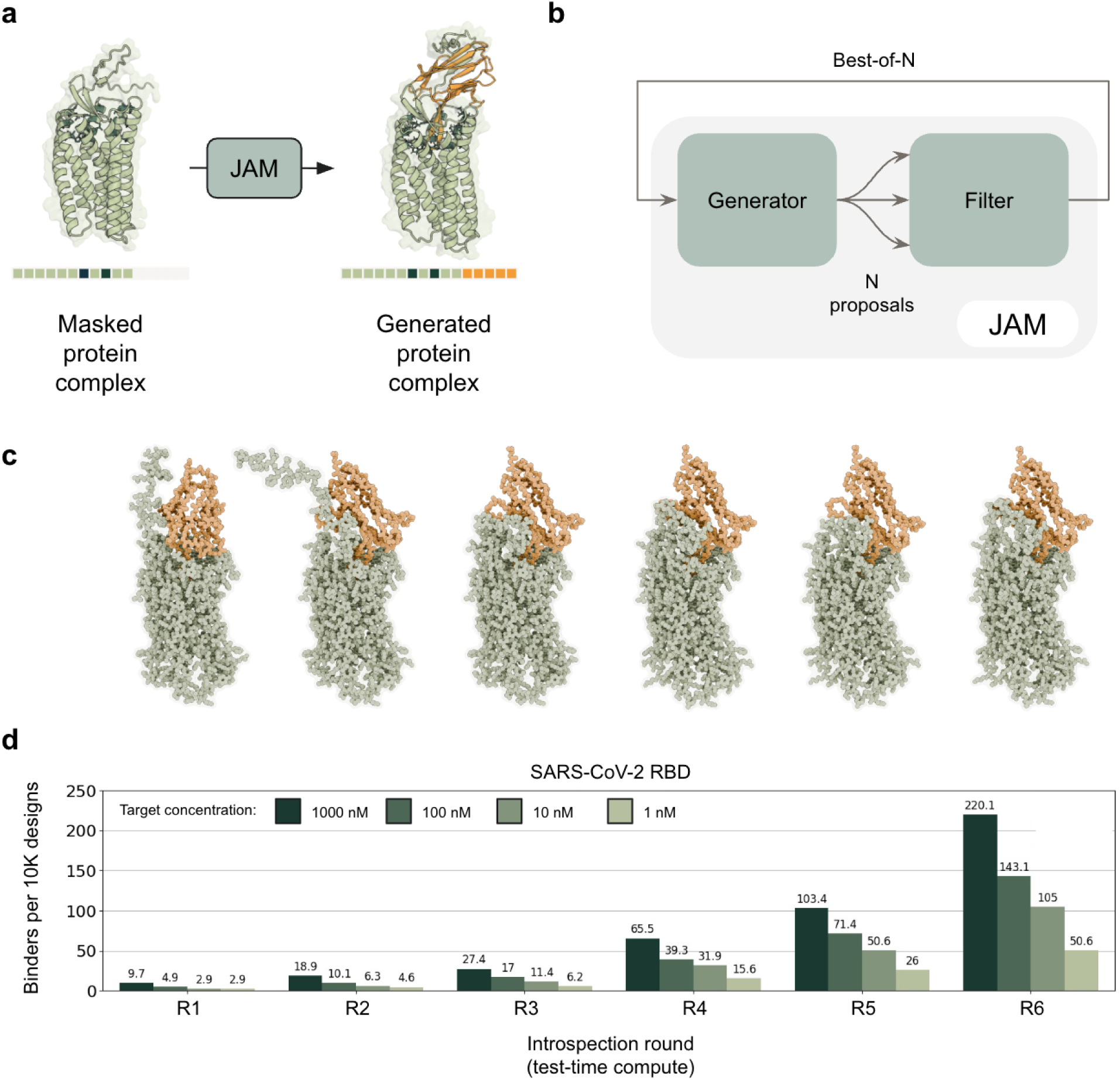
**a.** Illustrated example of JAM’s generative process for antibody design. Given partial target sequence and structure information, an epitope, and masked sequence and structure information for the antibody, JAM generates antibody sequence and structure. The sequence can be used to test the antibody experimentally. **b.** An introspection step involves generating *N* antibody-target complexes (sequence and structure) with JAM’s Generator module, followed by scoring and selecting top ranking complexes by the Filter module for re-prompting of JAM’s Generator. Filter-assigned scores are known to be predictive of antibody-target binding on internally collected datasets for targets unrelated to those explored in this work. **c.** Example of JAM-generated VHH-CXCR7 complexes over an introspection trajectory proceeding from Round 1 to Round 6. **d.** Number of anti-SARS-CoV-2 RBD binders observed (on-yeast) per 10,000 designs tested versus round of introspection and target concentration.

JAM contains a Generator and Filter module (Fig. 1b). The Generator is a generative model that creates complete protein complexes from partially specified ones as described above. The Filter module uses a complete protein complex as input and assigns a score predictive of the experimental viability of that complex. For multi-chain complexes (such as an antibody-target pair), this is equivalent to the likelihood of successful protein-protein interaction.

The Generator and Filter modules were pretrained on publicly available sequence and structure databases, the latter comprising both experimentally determined as well as model-predicted structures. Subsequently, during a post-training phase, all components of the system were aligned on internally generated data from over 70 design-build-test rounds to maximize the likelihood JAM generated complexes are experimentally viable. Each design-build-test round contains 10⁴-10⁵ JAM-generated complexes each paired with quantitative protein expression and binding measurements, and in aggregate (across rounds), spans a wide range of antibody-target combinations and single-chain proteins.

Previously, we observed that applying a single forward pass of JAM to generate VHHs *de novo* yielded nanomolar binders (measured on-yeast) at approximately 0.1% success rate. We hypothesized that generating high-quality antibodies may be too challenging in a single pass, and that giving JAM more opportunity to "think" by providing it the opportunity to refine its generated antibody-target complexes would improve success rates and affinities. Inspired by approaches in natural language processing for answering complex questions, such as Tree-of-Thoughts ^18^, we applied our fully-computational "introspection" approach where JAM’s Generator proposes multiple designs, the Filter scores them, and top-scoring designs are fed back into the Generator for iterative refinement of antibody sequence and structure along with target structure through resampling. Figure 1c illustrates an example introspection trajectory for a VHH-CXCR7 complex evolving over six rounds.

We previously demonstrated that three rounds of introspection improved on-yeast bind rates two-fold for SARS-CoV-2 RBD^17^. Here, extending to six total rounds before wet-lab testing, we observed superlinear improvements in bind rates and affinities as a function of introspection round. Six rounds (R6) produced nanomolar binder on-yeast at a rate of 2.2% — a 22-fold improvement over single-pass baseline (R1) (Fig. 1d). These results demonstrate that increased test-time computational resources enable JAM to achieve better results through enhanced reasoning capacity.

### Increased test-time compute improves *de novo* antibody design success rates **>60-fold for the GPCRs CXCR7 and CXCR4**

In our previous work, a single forward pass of the JAM pipeline targeting the GPCR CXCR7 yielded only one VHH binder from 20,000 designs tested. This singular design showed an affinity of 21.8 nM, but exhibited poor developability properties (Supp. Fig. 1)^17^. Encouraged by the superlinear performance gains observed with increased test-time compute on SARS-CoV-2 RBD, we hypothesized that similar application of increased test-time compute would transform rare successes on GPCRs into more robust hit-rates with improved affinities.

To test this hypothesis, we targeted CXCR7 with three rounds (R3) and six rounds (R6) of introspection. We also similarly targeted the closely-related GPCR, CXCR4, using three and six rounds of introspection. CXCR4 shares significant structural similarity with CXCR7, binding the same chemokine ligand (SDF1α)^1920^. This parallel targeting allowed us to not only assess the value of increased test-time resources, but also the generalizability of our design process, and evaluate the selectivity of our designs, a key requirement and hurdle for therapeutics targeting GPCRs in general^21^.

We generated 20,000 R3 *de novo* VHH designs for each CXCR4 and CXCR7, matching our previously-reported anti-CXCR7 R1 library in size. We also generated 107,355 R6 designs targeting CXCR7 and 89,671 R6 targeting CXCR4. Figure 2a illustrates the overall screening workflow used here to identify *de novo* VHH designs that bind the native GPCR, and Figure 2b provides two representative examples of JAM-generated VHH-GPCR complexes for both targets.

**Figure 2.**
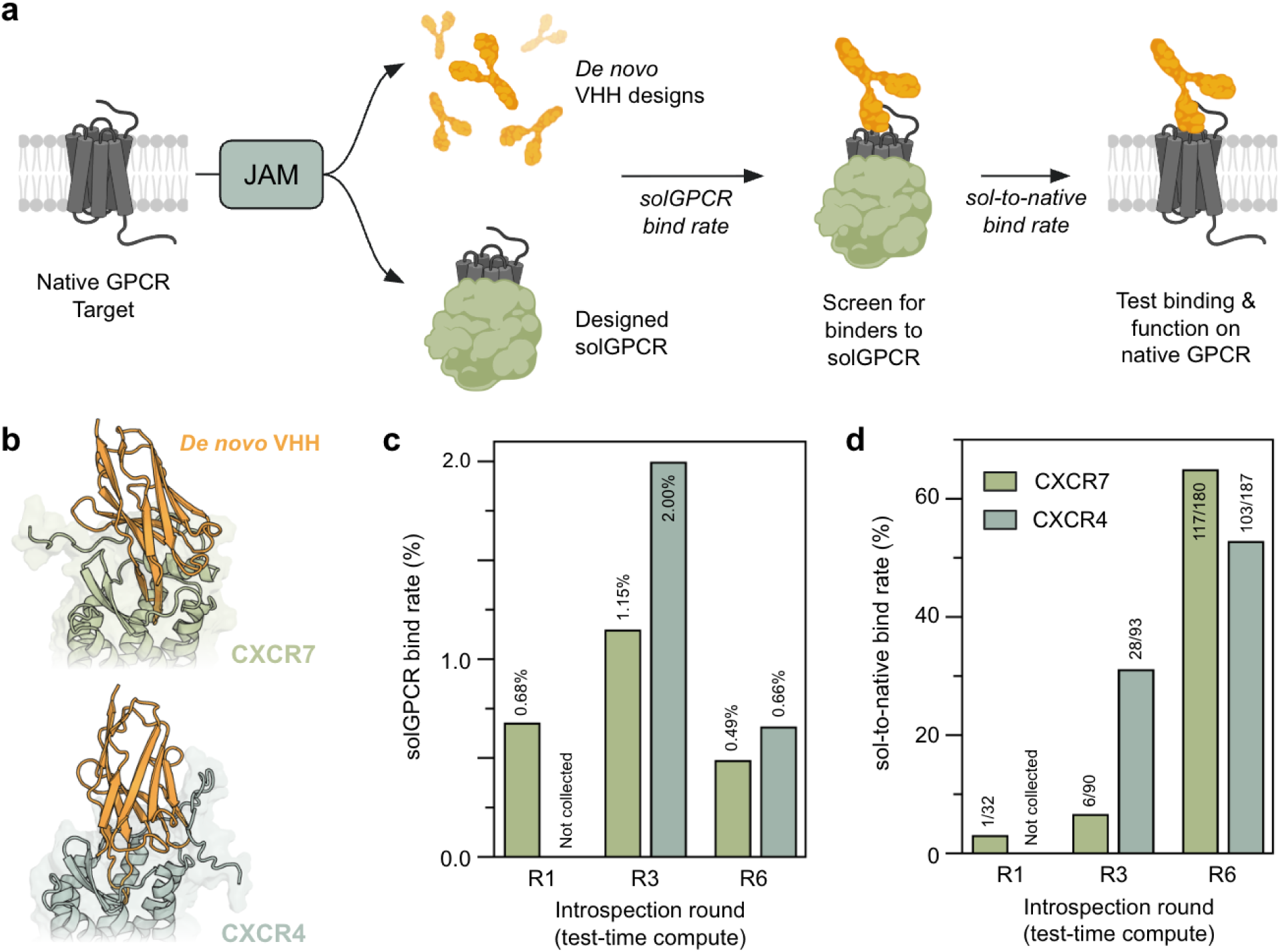
**a.** Given a GPCR target, JAM is used to generate both a soluble GPCR proxy (solGPCR) of the native GPCR as well as *de novo* VHH designs. The solGPCR is used as a proxy target to screen designed VHHs via yeast display. solGPCR binding VHHs are then tested on cells for binding to the native GPCR **b.** JAM-generated complexes of CXCR7 (light green) and CXCR4 (light blue) each bound by designed VHH antibodies (orange). The designed VHHs target the orthosteric binding site of the GPCR **c.** Bind rate by flow cytometry of *de novo* designs to solGPCRs as a function of introspection round. Observed % bind rate listed on top of each bar. **d.** Rate of solGPCR-binding designs that bind the native GPCR as a function of introspection round. Numbers on top of each bar shows the number of confirmed native GPCR binders / number of solGPCR binders produced recombinantly and tested on-cells.

To facilitate screening of these libraries, we utilized a JAM-designed soluble proxy of CXCR7 (solCXCR7) and CXCR4 (solCXCR4), made as described previously^17^. We refer to these collectively as solGPCRs. In these designed proxies, JAM replaces the transmembrane and intracellular region of the native multipass membrane protein with a stable, soluble scaffold while preserving critical extracellular structures of the native GPCR (Fig. 2a). We utilized the solCXCR7 from our previous work, which was 82% monomeric by size exclusion chromatography (SEC) and bound SDF1α with 8.1 nM affinity evaluated by biolayer interferometry (BLI). For CXCR4, we generated a solCXCR4 design, which was 73% monomeric, and bound SDF1α with 2.3 nM affinity (Supp. Fig. 2). These solCXCR4 and solCXCR7 designs were then used to screen the R3 and R6 *de novo* VHH libraries via yeast display for potential binders of the native GPCR (Fig. 2a, Supp. Fig. 3). A random subset of solGPCR binding designs were then reformatted as VHH-Fcs and expressed in ExpiCHO. The resulting clarified supernatants were tested in a single-point binding assay against the native GPCR at their maximal expression concentration, which ranged from 150 nM to 5 uM (Supp. Fig. 4, Methods). We refer to the percent of designs that bind the solGPCR proxy as “solGPCR bind rate” and the percent of solGPCR binders that bind the native GPCR as “sol-to-native bind rate” (Fig. 2a).

Interestingly, while we did not observe a correlation between test-time compute and solGPCR bind rate (Fig. 2c), we observed a strong correlation between test-time compute and sol-to-native bind rates for both GPCRs (Fig. 2d). For CXCR7, R6 designs demonstrated a 65% sol-to-native bind rate compared to just 3.1% and 6.7% for R1 and R3 designs, respectively (Fig. 2d). Similarly, for CXCR4 R6 sol-to-native bind rate was 55% for R6 designs compared to 30% for R3 (Fig. 2d). While we cannot calculate a true bind rate given all designs are first screened with the solGPCR, we estimate a lower bound of the bind rate to be 0.32% for both targets by multiplying the solGPCR bind rate by the sol-to-native bind rate (0.49% × 65% = 0.32% for CXCR7; 0.6% × 53% = 0.32% for CXCR4). This is a 64-fold increase in lower bound bind rate relative to a single round of introspection (1 binding design out of 20,000 tested, or 0.005%).

Taken together, these results demonstrate that increased test-time compute in the form of additional rounds of introspection significantly enhances the success rate of GPCR-targeting antibody designs.

### Top *de novo* designed antibodies are novel, demonstrate high selectivity, and low- to sub-nanomolar affinities, superior to that of clinical-stage molecules

Given the improved R6 hit rates relative to R3, we chose to focus characterization effort primarily on R6 designs (Supp. Fig. 5). We selected top designs from R6 pools by median fluorescence intensity (MFI) in the single point binding assay previously described, produced them at larger scale, and characterized their binding affinities to the native GPCRs on-cells (on-cell K_D_).

For CXCR4, among 35 R6 designs tested in VHH-Fc format, we observed a median on-cell K_D_ of 2 nM on the PathHunter-CXCR4 overexpression cell line, with negligible binding to the parent C2C12 cell line (Fig. 3a, Supp. Fig. 6). 7 of 35 designs demonstrated sub-nanomolar affinities (Fig. 3a). Notably, our top *de novo* designs showed affinities more than 10-fold stronger than that of Ulocuplumab, a fully human anti-CXCR4 monoclonal antibody developed by Bristol Myers Squibb that has undergone multiple Phase 1/2 clinical trials in hematologic malignancies (Fig. 3b)^22,23^. The highest affinity *de novo* design had a K_D_ of 370 pM (Fig. 3c).

**Figure 3.**
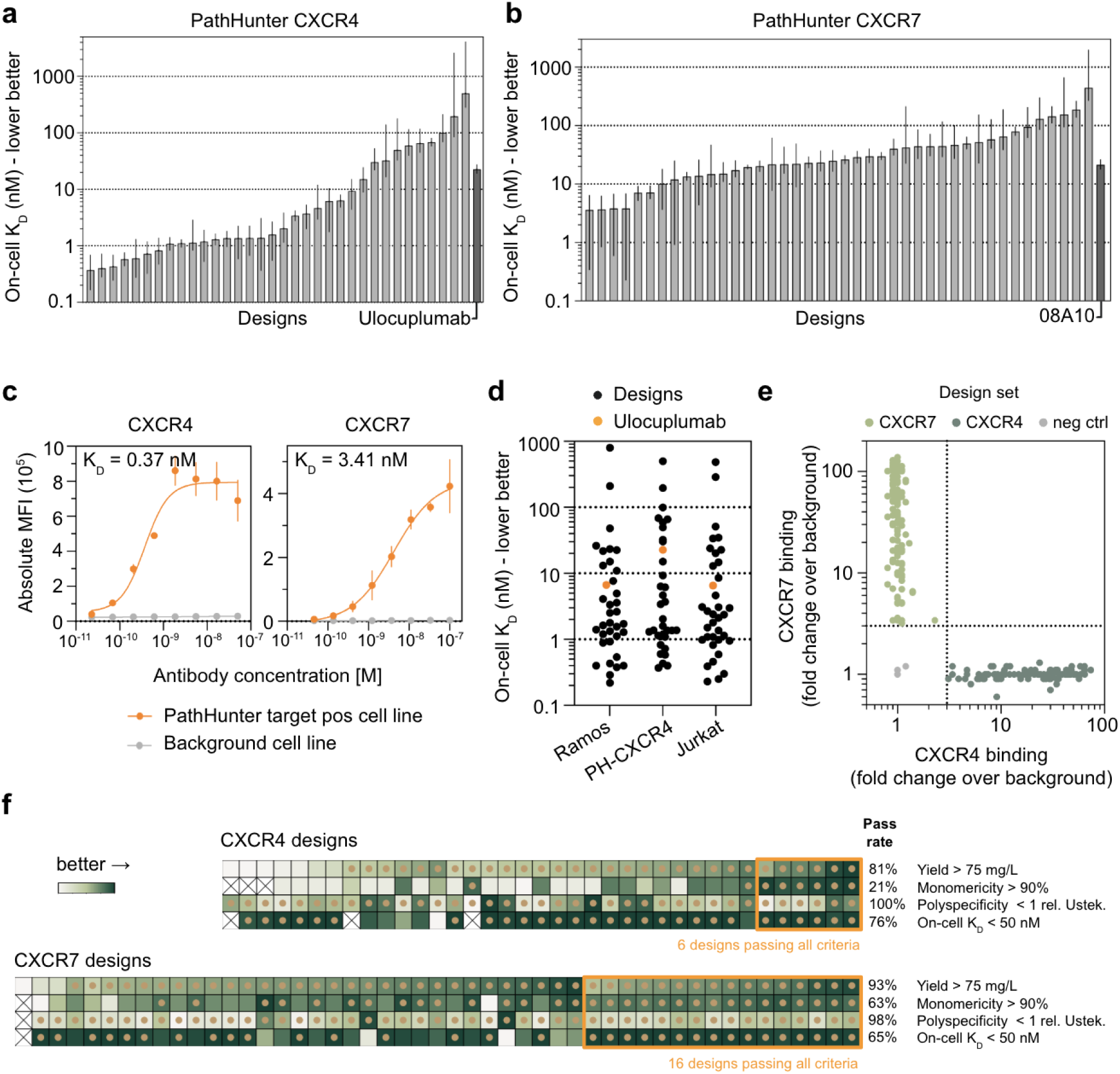
**a.** On-cell K_D_ measurements for titrations of anti-CXCR4 designs on PathHunter CXCR4 cell line. Error bars are 95% CI for bar plot and SD for binding trace. Weak binders for which an accurate on-cell KD was not calculable are excluded. Binding control is shown in dark grey. **b.** On-cell K_D_ measurements and titrations of anti-CXCR7 designs on PathHunter CXCR7 cell line as shown in panel (a). **c.** Binding trace of top anti-CXCR4 design (left) and anti-CXCR7 design (right) against their respective PathHunter target positive (orange) and background (grey) cell line. **d**. On-cell K_D_ for anti-CXCR4 designs against additional disease-relevant CXCR4 expressing cell lines (Ramos and Jurkat) **e.** Single-point binding signal for anti-CXCR4 and anti-CXCR7 at their highest expressed concentration against both CXCR4 and CXCR7 cell lines to evaluate on- and off-target binding. **f.** Developability heatmaps for CXCR4-targeting (top) and CXCR7-targeting (bottom) designs, showing production yield in ExpiCHO post 1-step purification, monomericity by SEC, and polyspecificity by BVP ELISA normalized to Ustekinumab. Designs passing the criteria are denoted by a brown dot. Designs meeting all developability criteria and an on-cell K_D_ threshold of <50 nM are highlighted by an orange box.

For CXCR7, we characterized 47 R6 designs as VHH-Fcs and found they had a median on-cell K_D_ of 34 nM on the PathHunter-CXCR7 overexpression cell line, with negligible binding to the parent C2C12 cell line (Fig. 3b, Supp. Fig. 6). Our top *de novo* designs bound CXCR7 with 2 nM on-cell K_D_ (Fig. 3c). This is a >10-fold improvement over the single CXCR7-binding *de novo* VHH we reported previously, which was generated with a single forward pass of JAM^17^.

As an additional point of comparison, we characterized the affinity of a patented CXCR7 VHH-Fc, 08A10. 08A10 was developed by Ablynx in March 2012, and discovered through a llama immunization with cells engineered to overexpress CXCR7, followed by multiple rounds of screening and counter-screening of the B-cell repertoire via phage display against target positive and target negative cells^22^. This represents a gold-standard screening cascade, which nonetheless often fails for GPCR targets. Our top CXCR7-targeting designs exhibited nearly a 10-fold higher affinity compared to this benchmark which showed an on-cell K_D_ of 18 nM when assayed concurrently (Fig. 3b).

To further validate the specificity and robustness of our designs, we tested binding on disease-relevant Ramos and Jurkat cell lines, which express varying levels of CXCR4 and provide orthogonal cellular backgrounds and surface receptor copy numbers (Supp. Fig. 7)^24^. These cell lines are widely used preclinical models for B- and T-cell malignancies, where CXCR4 mediates critical roles in cell trafficking, survival, and microenvironmental retention^17,25^. Our designs showed similar on-cell K_D_ values across these different cell types, providing additional confidence that they specifically recognize CXCR4 across diverse cellular contexts (Fig. 3d, Supp. Fig. 8).

Since CXCR4 and CXCR7 share a common endogenous ligand, SDF1α, we questioned whether our *de novo* designed binders, which generally target the SDF1α binding site, showed undesirable off-target binding. To assess specificity, we cross-tested all binders of native CXCR4 and CXCR7 against the alternate GPCR target. We observed no cross-reactivity, confirming that our designs were specific despite the significant structural similarity between these receptors (Fig. 3e).

To evaluate the early-stage developability of our lead designs, we assessed production yield in ExpiCHO, monomeric purity by SEC upon a high-throughput Protein A-based purification and without additional polishing workflows, and polyspecificity using a BVP ELISA assay (Methods, Supp. Fig. 9-10). To classify a design as having favorable affinities and early stage developability properties, we defined “passing” criteria for each assay as follows: on-cell K_D_ < 50 nM, yield > 75 mg/L, monomericity > 90%, and a BVP score lower than that of Ustekinumab, an approved monoclonal antibody which is moderately polyreactive^26^.

6/37 (16%) and 16/49 (33%) of CXCR4- and CXCR7-targeting designs, respectively, met all four criteria (Fig. 3f). We additionally quantified each design’s humanness as its percent sequence identity to the nearest human V-gene germline sequence - a metric relevant for immunogenicity risk assessment. The majority of designs showed humanness scores with >75% identity to human germline VH, comparable to humanized clinical antibodies and an improvement upon our previous work (Supp. Fig. 11)^17^.

JAM-generated VHHs had novel sequences distinct from those in existing sequence databases. To assess sequence novelty, we compared designs against all sequences in the NR (NCBI)^27^, OAS Unpaired (OPIG)^28^, and INDI (NaturalAntibody)^29^ databases, collectively containing over 3 billion sequences. All JAM-designed VHHs targeting both CXCR4 and CXCR7 showed greater than 18% sequence dissimilarity from their closest database matches (Supp. Fig. 12a).

Additionally, the designed VHH-GPCR complexes were structurally novel in terms of the predicted binding pose of the VHH. We assessed structural novelty by comparing JAM-generated VHH-target complexes against all antibody-antigen structures in SAbDab (OPIG)^30^. For each experimentally validated JAM complex, we identified the most similar known structure by aligning on target chains and calculating alpha-carbon RMSD between the antibody components, which we refer to as “target-aligned antibody RMSD” (Methods). All designs exhibited target-aligned antibody RMSD values exceeding 7 Å from the nearest SAbDab structure, with median RMSDs of 10 Å and 12.2 Å for CXCR4- and CXCR7-targeting designs, respectively (Supp. Fig. 12b). Similar analysis of just the CDR regions and CDR3 alone showed all designs maintained greater than 6 Å RMSD in these regions, with median values of 9.8 Å and 14 Å for CXCR4- and CXCR7-targeting designs, respectively (Supp. Fig. 12b).

These results demonstrate that purely *de novo*, we can generate novel single-domain antibody constructs with affinities matching or exceeding those of clinical-stage antibodies, with a substantially higher hit-rate than traditional approaches such as immunization and panning.

### Top *de novo* designed antibodies are low- to sub-nanomolar antagonists, and a subset are first-in-class agonists of CXCR7

All JAM-generated *de novo* VHH designs against CXCR7 and CXCR4 target extracellular residues on the main 7-transmembrane (7TM) barrel of the GPCR and none exclusively contact the N-terminal loop. For most designs, the CDR3 loop extends into the orthosteric pocket of the GPCR and spatially overlaps with regions critical for SDF1α engagement (Fig. 4a). We therefore hypothesized that these designs would be functional antagonists, blocking SDF1α binding and/or SDF1α-mediated signaling For CXCR4, therapeutic efforts have largely focused on antagonizing its interaction with the chemokine SDF1α^25^. We therefore evaluated the ability of our designs to outcompete SDF1α binding to CXCR4 on the disease-relevant Ramos and Jurkat cell lines (Methods). Of 37 R6 *de novo* designs tested, 34 demonstrated competitive displacement of SDF1α from CXCR4 on both Ramos and Jurkat cell lines (Fig. 4b, Supp. Fig. 13). Consistent with their on-cell KD values, top designs showed picomolar IC50s and were substantially stronger competitors than the clinical-stage monoclonal antibody Ulocuplumab (Fig. 4b,c).

**Figure 4.**
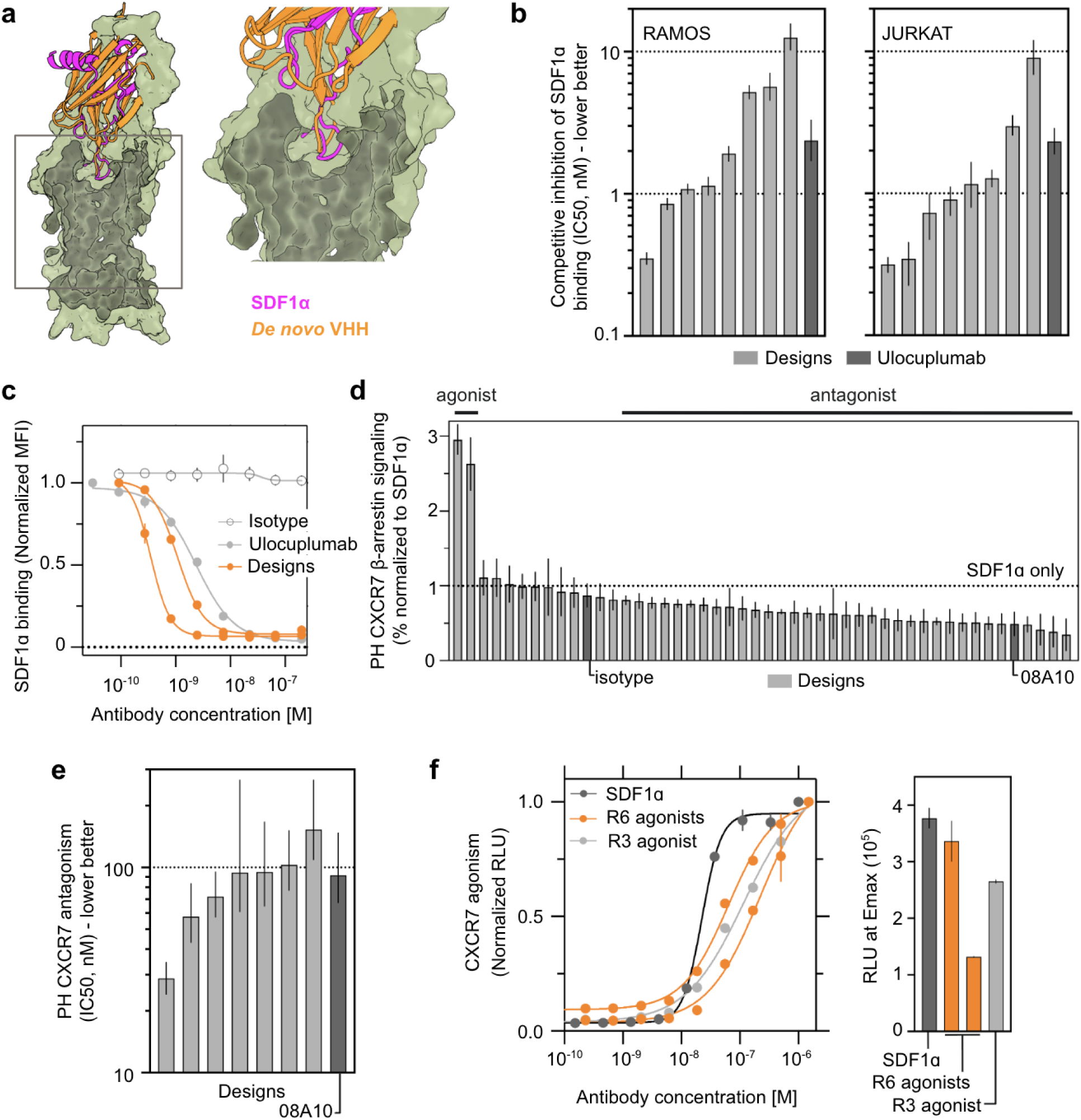
**a.** Structural models show the predicted binding mode of agonist designs on CXCR7 (green); magenta = SDF1α, orange = JAM-designed VHH. **b.** Competitive inhibition of SDF1α binding to native CXCR4+ cells (Ramos, Jurkat) by *de novo* designs and control antibody, Ulocuplumab. Bars represent IC50 values; error bars indicate 95% CI. **c.** Flow cytometry-based titrations show displacement of fluorescently labeled SDF1α from Ramos cells by designs (orange), Ulocuplumab (light grey) and an isotype control (grey, hollow symbol). **d.** β-arrestin signaling on PathHunter CXCR7 cells in response to antibody designs added at 200 nM in conjunction with SDF1α, normalized to SDF1α-only. Designs span both agonists and antagonists. Isotype and positive control (pos ctrl) are labeled accordingly **e.** Antagonistic potency (IC50) of anti-CXCR7 designs and control (08A10) against SDF1α-induced β-arrestin signaling. Error bars indicate 95% CI. **e**. EC50 agonism (β-arrestin signaling) curves of the two agonistic R6 designs targeting CXCR7 (left); right: comparison of Emax (RLU) with SDF1α (dark grey) and R3 agonist (light grey).

To evaluate CXCR7-targeting designs, we characterized the same 49 R6 *de novo* designs profiled in our affinity studies for their ability to modulate SDF1α-mediated β-arrestin signaling (Methods). In a single-point assay, 200 nM antibody was incubated with the PathHunter CXCR7 cell line prior to addition SDF1α added at 30 nM (Methods). We observed 35 out of 47 (74%) designs were antagonistic, with top designs showing 75% reduction in SDF1α-mediated signaling, exceeding that of the benchmark molecule 08A10 (Fig. 4d). The top 7 designs were further characterized with 8-pt titrations and the top among these showed IC50s of 29 and 58 nM (Fig. 4e, Supp. Fig. 14).

Strikingly, in the same assay, two *de novo* designs were identified as agonists, increasing the observed β-arrestin signaling in the described single-point assay (Fig. 4d). These designs were both confirmed to independently activate β-arrestin signaling in further characterizations, with our top agonist design showing an EC50 of 63 nM and similar Emax to the natural ligand agonist SDF1α, which had an EC50 of 22.5 nM when measured concurrently (Fig. 4f). To the best of our knowledge, these represent the first antibody agonists reported for this biologically central GPCR, let alone agonists that are designed fully computationally.

We also observed that agonist activity was not exclusive to R6 designs. We identified an R3 *de novo* design with agonism average EC50 of 108 nM across experimental batches. 3 of 5 remaining R3 designs were antagonistic, though these were not characterized with the same degree of replication (Supp. Fig. 15).

These results demonstrate that our *de novo* designed antibodies not only bind with high affinity to their targets, but also modulate receptor function in therapeutically relevant ways and with therapeutic-grade potencies.

### Given a single data point, experiment-guided steering of JAM yields >300 diverse CXCR7 agonists with improved function and developability

Prior work in language modeling has shown that high-capacity large language models (LLMs) can “learn” from new data without updating their weights or training additional models. Instead, new information can be incorporated directly into the prompt, enabling the model to infer patterns at test-time and generate appropriate, task-specific outputs. This approach, known as in-context learning, assumes that the LLM already contains latent knowledge acquired during pre- and post-training, and that prompting serves to activate and steer that knowledge toward the task at hand^31^.

In protein design, recent work has shown that prompting protein language models or inverse folding models with the sequence and structure of an antibody known to bind a target, and asking them to propose mutations, can yield variants with improved affinity at double-digit percent success rates, without requiring additional training on application-specific experimental data^32,33^. While these studies explored only a narrow set of mutations around the input sequence and required an experimentally determined antibody structure, Hayes et al. (2024) showed that ESM3 can generate novel fluorescent proteins highly dissimilar to natural ones when prompted with just the chromophore sequence and structure. Although the best initial design exhibited modest fluorescence, it could be then used to re-prompt ESM3, leading to a second-round design with brightness comparable to natural fluorescent proteins^9^.

These results point to a larger emerging paradigm of **experiment-guided steering** of protein generative models for protein design. Highly data-efficient and lightweight, this framework uses a small number of experimentally validated proteins with desirable properties to simply (re-)prompt a protein generative model at test-time to produce new proteins that are semantically “like” the prompted examples, some of which may exhibit even stronger or improved characteristics (Fig. 5a).

**Figure 5.**
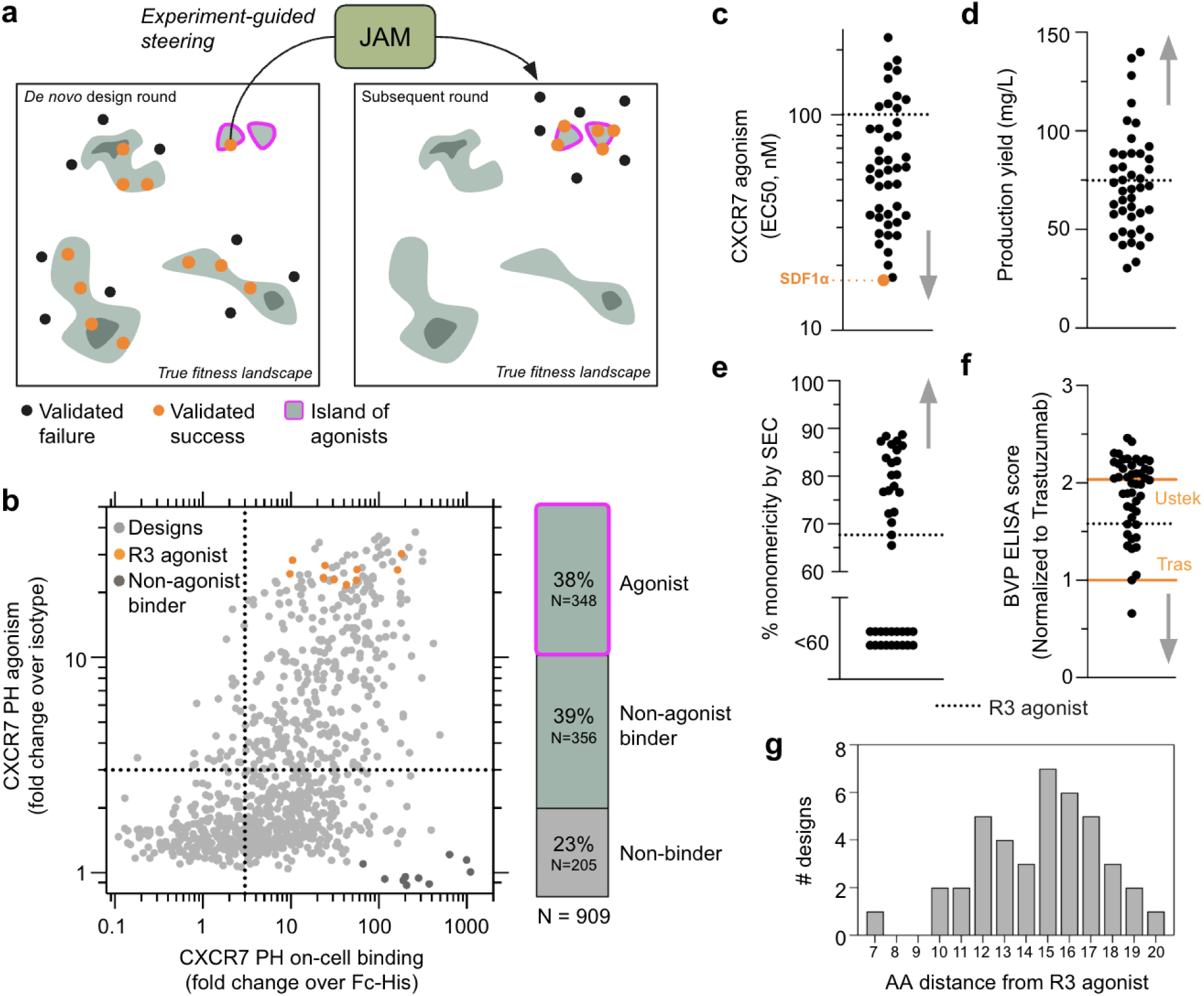
**a.** Cartoon illustration of experiment--guided steering of JAM. In an initial *de novo* design round JAM produces designs that successfully bind (orange) or do not (black). A small “island” (magenta border) in the fitness landscape is populated not just by binders, but by binder-agonists, and JAM successfully proposes a design in that region. The JAM-generated complex (sequence + structure of designed antibody + target) of that single agonist is used to re-prompt JAM to generate more designs that are semantically similar. These are experimentally characterized in a subsequent round, and more agonists with diverse properties in other dimensions are identified. **b.** 909 designs evaluated for CXCR7 on-cell binding (x-axis) and agonism via β-arrestin signaling (y-axis). Dotted lines indicate three-fold over background (isotype or Fc-His) used as classification thresholds for binding and agonism. The R3 agonist (orange) and a non-agonist binder (08A10, dark gray) were also evaluated. For panels **c-f**, a grey arrow indicates the direction of more favorable properties and the dotted line represents the R3 agonist produced and characterized concurrently **c**. Agonism EC50 values for designs assayed for CXCR7 β-arrestin signaling using the PathHunter CXCR7 cell line. SDF1α (orange dot) was also assayed. **d.** Purification yield of VHH-Fc binders from 24-well ExpiCHO production **e.** % monomericity post-purification as determined by Size Exclusion Chromatography. **f.** Polyspecificity scores as measured by BVP ELISA and normalized to <u>Tras</u>tuzumab (lower orange line at y=1). <u>Ustek</u>inumab (upper orange line) was assayed in the same plate **g.** Histogram showing the number of JAM-generated designs as a function of their distance in amino acid mutations from the R3 agonist sequence.

Encouraged by our discovery of CXCR7 agonists, we wondered if we could use one of them to steer JAM to generate more agonists with improved functional and developability properties. We chose to focus on the R3 agonist due to its strong functional activity, but poor monomericity. To generate more agonist candidates we took the JAM-generated VHH-CXCR7 complex corresponding to this agonist and simply re-prompted JAM with it to produce new VHH designs, avoiding any context-specific training of JAM or post-hoc models.

In two weeks, including re-prompting JAM, we characterized the 909 designs in a single-point screen for agonism at each antibody’s highest expressed concentration (Fig. 5b). Remarkably, we observed that 704/909 designs showed binding to CXCR7, and of these, 348 demonstrated agonist activity at least three-fold greater than isotype controls and 08A10 (a non-agonizing binder) — a >7500-fold enrichment compared to our initial discovery rate in R6.

For detailed characterization, we selected 43 designs, prioritizing those with higher single-point agonism signal at 30 nM antibody concentration, followed by those with the highest supernatant expression titer, subject to agonism signal >0.8 that of the R3 agonist parent measured in the same experiment. 34 of the 43 designs proved to be stronger agonists than our original hit, with the top design exhibiting an agonism EC50 equivalent to that of CXCR7’s natural ligand SDF1α

(Fig. 5c). Additionally, approximately half of the designs showed improved production yields while maintaining polyspecificity scores below that of the clinical benchmark Ustekinumab. About half of the designs also demonstrated significantly improved monomericity, though none exceeded 90% monomericity in our high-throughput purification protocol (Fig. 5d-f), suggesting some developability challenges may remain. Importantly, these JAM-generated designs were not trivial variations of the R3 agonist. Though they were not selected to be diverse with respect to the R3 agonist parent, they contained between 5 and 19 mutations relative to the R3 agonist across the full-length sequence, and between 0 and 11 mutations in the CDRs, indicating substantial diversity in the proposed solutions (Fig. 5g).

This approach demonstrates a powerful method for incorporating extremely limited experimental data (N=1) in a straightforward manner, without requiring model retraining or custom model development. By prompting JAM with a single successful example, we transformed what was initially a rare discovery into a robust and diverse lead series of functional CXCR7 agonists with substantially improved developability.

## Discussion

We demonstrated that test-time scaling dramatically enhances *de novo* antibody design against SARS-CoV-2 RBD and GPCRs CXCR4 and CXCR7. By allowing JAM to iteratively introspect on its outputs for six, entirely computational rounds, we increased bind rates 20-70x and obtained dozens of single-digit nanomolar or picomolar binders — in some cases with affinities >10-fold stronger than benchmark molecules discovered through conventional approaches like immunization and phage display. Our designs exhibited high target selectivity despite CXCR4 and CXCR7 being structurally similar enough to bind the same ligand SDF1α. Most of our top designs were potent antagonists, effectively competing with SDF1α for GPCR binding or inhibiting receptor signaling.

Strikingly, we also discovered three *de novo* designed VHHs that agonized CXCR7 — to our knowledge, the first computationally designed antibody agonists of any GPCR and the first antibody agonists of CXCR7. While antibody agonists are known to be exceedingly rare, we demonstrated through experiment-guided steering that they can be efficiently generated given a single experimental datapoint as an initial bearing. By simply re-prompting JAM with the original JAM-generated complex of a successful *de novo* agonist, we produced over three hundred diverse agonists in just weeks, with top designs showing improved developability and potency rivaling SDF1α — a natural ligand whose interaction with CXCR7 has been developed and refined by evolution over 400 million years^34^. This efficient integration of experimental data highlights how test-time techniques can be applied not only for *de novo* design, but also for incorporating experimental feedback with minimal data (N=1) without requiring application-specific training.

Limitations of our study include its focus on two related GPCRs, leaving generalizability to other GPCR families and multipass membrane proteins to be determined. Additionally, while we improved JAM’s ability to generate more human-like sequences compared to our previous work, our designs still possess humanness comparable to humanized rather than fully human clinical-stage antibodies. We also focused exclusively on VHH formats for wet lab experimental simplicity, though based on our previous work demonstrating JAM’s capability with paired VH+VL formats, we expect these test-time scaling benefits to extend there as well.

Ongoing work is exploring the applicability of these results to other challenging targets, including ion channels and transporters. We are also investigating the potential for further gains with additional rounds of introspection, as we have not yet observed a plateau in performance improvement.

Our results have significant implications for therapeutic discovery against multipass membrane proteins. These targets, comprising approximately two-thirds of cell surface proteins, offer numerous possibilities to influence disease biology but are targeted by less than 10% of current biologics. This limited success stems from multiple challenges: restricted extracellular epitopes, high sequence similarity between family members, difficulties in using membrane-bound proteins as screening reagents, and sometimes, the need to target specific conformational states. *De novo* antibody design with test-time scaling may help to address these challenges by enabling atomic-precision engineering of binding interfaces.

The improved hit rates achieved through test-time scaling now enable direct screening against native GPCRs on cells, eliminating the need for soluble proxies that introduce unknown false negative rates. Direct screening would not only shorten timelines, but also allow evaluation of binding and function in settings with higher therapeutic relevance. As hit rates continue to improve, we anticipate being able to design smaller sets of antibodies with high confidence of success, eventually enabling performance assessment in high relevance settings like patient primary cells or even directly in animal models.

We believe that expanding the reasoning capacity of biomolecular generative models through increased test-time computation will become a fundamental "scaling law" important for the design of biological systems. Just as test-time reasoning is rapidly transforming language model capabilities and enabling machines to solve increasingly complex problems, test-time scaling in biological design may too soon follow a similar trajectory. This emerging paradigm offers a path toward designing therapeutics for previously intractable targets, and we believe will be critical to advancing AI-guided drug discovery.

## Experimental Methods

### *De novo* designed yeast surface display library construction

To construct the yeast surface display libraries, oligonucleotides encoding the designed VHH antibodies were ordered from Twist Biosciences as 300nt oligo pools with flanking BsaI recognition sites for Golden Gate assembly. All DNA was codon-optimized for expression in S. cerevisiae. Golden gate assembly reactions were run overnight using PCR amplified oligo pool DNA to clone into the yeast display vector, pCTcon2.

Purified golden gate reactions were electroporated into NEB 10-beta electrocompetent E. coli (New England Biolabs) using the pre-set bacterial protocol on the Gene Pulser Xcell Electroporation System (BioRad). Serial dilutions of the bacterial transformants were plated and verified to represent a greater than 100-fold coverage of the library. The resulting plasmid library was extracted from the bacterial cultures using a QIAprep Spin Miniprep kit (QIAGEN).

The assembled libraries were linearized and transformed into S. cerevisiae strain EBY100 (ATCC) using a standard lithium acetate and DTT-based yeast electroporation protocol as described by Van Deventer et al.^35^. Transformants were serial diluted and plated post recovery to verify library coverage was at least 100-fold. Yeast transformants were cultivated in synthetic dextrose medium with casamino acids (SDCAA) pH 4.5 (Teknova) shaking at 30°C overnight.

### Cell sorting of yeast surface displayed antibody libraries

Yeast libraries were grown to saturation overnight in SDCAA pH 4.5 media (Teknova) shaking at 30°C. Each library was passaged into fresh SDCAA pH 4.5 media at a 25X dilution and grown for 2-4 hours before pelleting via centrifugation at 2000 x g for 5 minutes. To induce the libraries, cell pellets were resuspended to a OD600 of 1 in synthetic galactose medium with casamino acids (SGCAA) (Teknova) and incubated at 20°C for 20 hours.

Fluorescence-Activated Cell Sorting (FACS):

For RBD data, SARS-CoV-2 (COVID-19) S protein RBD (AA319-537) with a C-terminal His Tag (SPD-C52H3; AcroBiosystems) was used for sorting. A first FACS enrichment was performed at 1000 nM. This enriched population was then reinduced and sorted for a second enrichment at four antigen concentrations (1000, 100, 10, 1 nM) to isolate binders with various affinities (Table 1).

**Table 1:**
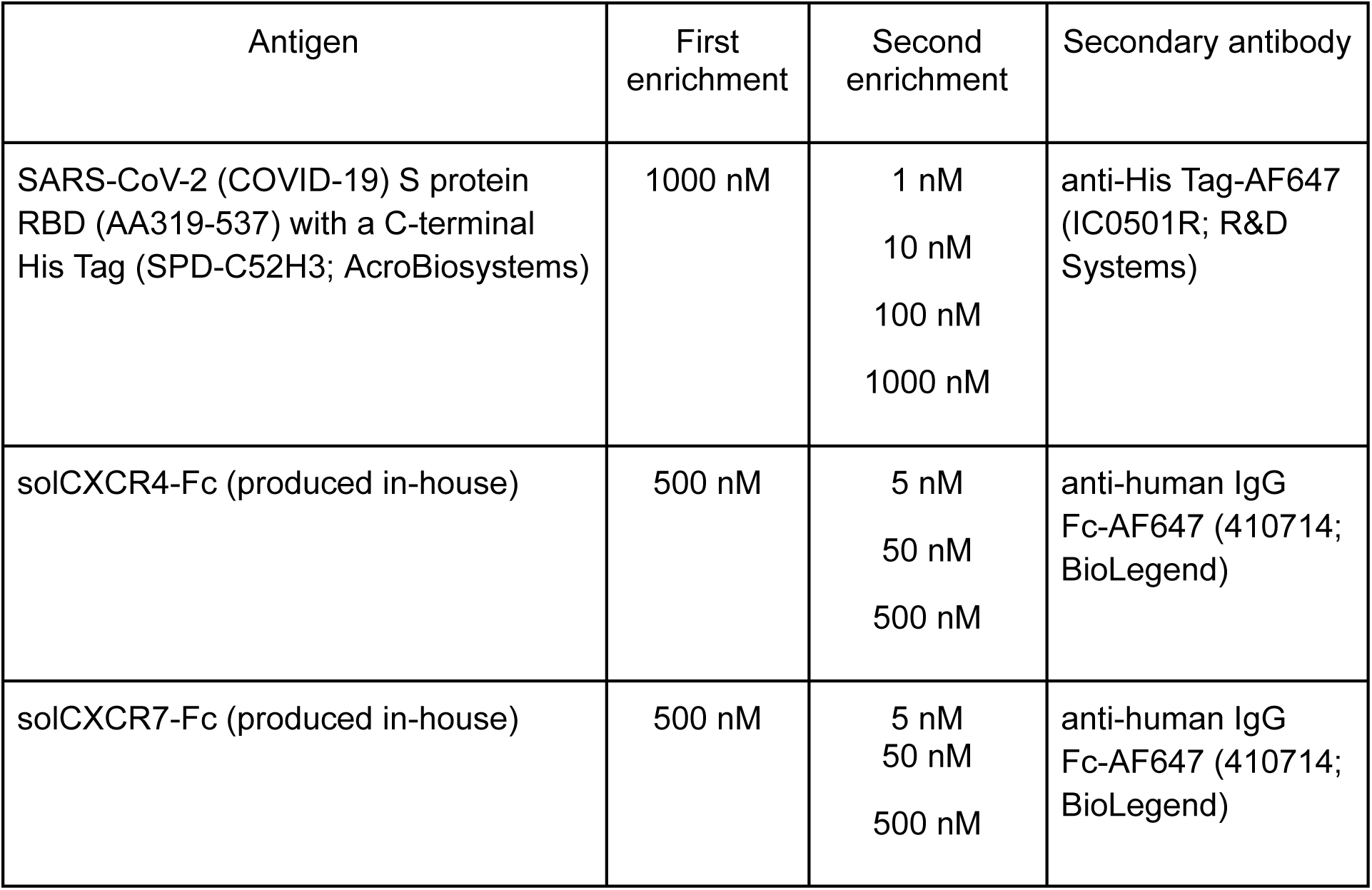
Yeast surface display library enrichments: antigen concentrations and reagents.

| Antigen | First enrichment | Second enrichment | Secondary antibody |
| --- | --- | --- | --- |
| SARS-CoV-2 (COVID-19) S protein RBD (AA319-537) with a C-terminal His Tag (SPD-C52H3; AcroBiosystems) | 1000 nM | 1 nM<br>10 nM<br>100 nM<br>1000 nM | anti-His Tag-AF647 (IC0501R; R&D Systems) |
| solCXCR4-Fc (produced in-house) | 500 nM | 5 nM<br>50 nM<br>500 nM | anti-human IgG Fc-AF647 (410714; BioLegend) |
| solCXCR7-Fc (produced in-house) | 500 nM | 5 nM<br>50 nM<br>500 nM | anti-human IgG Fc-AF647 (410714; BioLegend) |

For GPCRs, Fc-fusions of each solGPCR (solCXCR4-Fc and solCXCR7-Fc) were produced and purified in-house according to methods described in the mammalian protein production and purification sections and used for sorting. A first FACS enrichment was performed with 500 nM Fc-tagged solGPCR. This enriched population was then reinduced and sorted for a second enrichment at three antigen concentrations (5 nM, 50 nM, 500 nM) to isolate binders with various affinities (Table 1).

For sorting, induced yeast cells were washed twice with 1% PBSA (1% BSA in 1X PBS). The libraries were incubated with anti-c-Myc-AF488 antibody (1:100x dilution, 16-308, Sigma-Aldrich) to label yeast cells displaying full length VHHs, and the desired antigen concentrations to evaluate antigen binding for 1 hour at room temperature. The samples were spun down at 4°C and washed twice with ice-cold 1% PBSA to remove unbound antigen and c-Myc antibody. 1:100X dilution of the relevant secondary antibody (Table 1) was added and the samples were incubated on ice for 30 minutes. The samples were spun down at 4°C and washed twice with ice-cold 1% PBSA to remove any excess secondary antibody.

Samples were sorted on a Sony SH800 cell sorter using normal or ultra purity mode depending on the stage of enrichment. Sorted binders were collected in SDCAA pH 4.5 media and shaken at 30°C for 2-3 days until saturated.

### Next generation sequencing of sorted yeast surface display populations

Binders isolated from the second enrichment were yeast miniprepped using a Zymoprep Yeast Plasmid Miniprep II kit (Zymo Research). DNA was then electroporated into NEB 10-beta electrocompetent E. coli (New England Biolabs) and recovered in LB liquid medium supplemented with carbenicillin. E.coli cultures were bacterial miniprepped using a QIAprep Spin Miniprep kit (QIAGEN) and then submitted for sequencing using Oxford Nanopore technology.

For YSD data, nanopore reads were mapped to the coding DNA of JAM-generated designs, and the number of reads that aligned to each design are tabulated. Due to sequencing noise, some designs have non-zero but low read count levels, whereas designs truly present in the binding population have higher read counts. To classify binders from non-binders, we set a read count threshold based on the empirical cumulative distribution function (eCDF) of read counts in the sample. Typically, 80-90% of reads are accounted for by a small number of designs, and the presence of clear “elbow” in the eCDF suggests a natural read count threshold that separates non-binders from binders. Empirically, we have validated this strategy identifies binding designs such that when expressed recombinantly and tested individually, designs are highly likely to bind the target.

### Mammalian protein production

Genes for recombinant protein production were synthesized either as gene fragments (Integrated DNA Technologies) for selected YSD binders or as oligo pools (Twist Biosciences) for random colony picking. All constructs were codon-optimized for expression in Homo sapiens and cloned into pcDNA3.4 (Invitrogen) using Golden Gate cloning as described previously^17^. All in-house produced antibodies and Fc-tagged proteins are of human IgG1 antibody subclass and contain the L234A, L235A, P329G (LALA PG) mutations to reduce effector function^36^. VHH-Fcs were produced in a VHH-G4S-Fc format.

For agonist designs developed through reprompting of JAM, an oligo pool of 6000 unique designs was utilized. Briefly, purified golden gate reactions were electroporated into NEB 10-beta electrocompetent E. coli (New England Biolabs) using the pre-set bacterial protocol on the Gene Pulser Xcell Electroporation System (BioRad). Dilutions of the bacterial transformants were plated on LB agar plates containing carbenicillin for random colony picking of 909 clones.

Colonies were inoculated into 96 deep well blocks containing LB or Terrific Broth (TB) media supplemented with carbenicillin and grown overnight until saturation. Pelleted cultures were then miniprepped to obtain transfection-grade plasmid.

For protein production, plasmids were transiently transfected into ExpiCHO cells (1 μg DNA/ mL cell culture) using the ExpiCHO Expression System (Gibco). Following harvest 6 days after transfection, the cell culture supernatant was clarified by centrifugation at 2000-3000 x g for 20-30 minutes. Concentrated (10X) phosphate buffered saline (PBS), pH 7.4 was added to the supernatant to achieve a final concentration of 1X PBS.

### Protein purification

Fc-containing proteins were purified using either rProtein A Sepharose Fast Flow antibody purification resin (Cytiva) or Pierce™ Protein A/G Magnetic Agarose Beads (Thermo Scientific). The resin/beads were equilibrated in 1X PBS then added to the ExpiCHO supernatant and mixed for 1 hour to achieve binding. Two to three washes with 1X PBS were performed to remove residual supernatant and non-specific proteins. Bound proteins were then eluted with IgG Elution Buffer, pH 2.8 (Thermo Scientific), then neutralized with 1 M Tris-HCl (Invitrogen) to approximately pH 7.

Following purification, the eluted protein solutions were buffer exchanged into PBS using Zeba™ Spin Desalting Columns or Plates (Thermo Scientific). The final protein concentration was quantified via A280 and a production yield is determined by dividing the resultant amount of purified protein by the production volume.

### Monomericity - Size Exclusion Chromatography

Following one-step Protein A bead-based purification, proteins were analyzed on an SRT C SEC 300 (235300-7830; Sepax Technologies). The column was pre-equilibrated in 4 column volumes of the mobile phase buffer (150 mM sodium phosphate at pH 7.0) before the first injection. For each run, 75 μL (containing ∼17 μg of each sample) was injected into the pre-equilibrated column using an Agilent 1100 system, equipped with an autosampler, at a flow rate of 1 mL/min. The retention time for each sample was determined based on the major peak at 280 nm.

### Polyspecificity - Baculovirus Particle (BVP) ELISA

BVP ELISA was performed as described in Jain et al.^26^. 50 μL of 1% baculovirus particles in PBS (Medna scientific) were diluted with equal volume of 50 mM sodium carbonate (pH 9.6) per well, and incubated on high bind ELISA plates (3369; Corning) at 4°C overnight with shaking. The next day, unbound BVPs were removed from the wells and the plate was washed 3x with 100 μL of 1X PBS. The plate was inverted on a Kimwipe on the bench to ensure the wells dried completely. All remaining steps were performed at room temperature. 100 μL of blocking buffer (PBS with 0.5% BSA) was added to the plate and incubated for 1 hour at room temperature with shaking at 450 rpm. Following incubation, the plate was washed 3x with 100 μL of 1X PBS. Next, antibodies were diluted to 1 μM in blocking buffer and wells were treated with 50 μL for 1 hour at room temperature with shaking at 450 rpm. The plate was then washed 6x with 100 μL of 1X PBS. 50 μL of 10 ng/mL secondary anti-human IgG-HRP conjugate (Jackson ImmunoResearch) was added to the wells and incubated for 1 hour at room temperature. The plate was then washed 6x with 100 μL of 1X PBS. Finally, 50 μL of room temperature TMB substrate (34021; Fisher Scientific) was added to each well and incubated for ∼5-10 minutes with gentle shaking. The reactions were stopped by addition of 50 μL 4N sulfuric acid to each well.

Absorbance at 450 nm was read on an iD3 SpectraMax Microplate Reader (Molecular Devices). BVP scores were determined by normalizing raw absorbance to control wells with no test antibody (i.e. BVP-only control), and normalized to Trastuzumab (ICH-4031; Ichor Bio) or Ustekinumab (HY-P9909; MedChem Express) scores assayed concurrently. Data is presented as the average of two replicate wells.

### Cell binding assays via flow cytometry

CHO-K1 (ATCC), PathHunter CHO-K1 CXCR7 β-arrestin (DiscoverX), C2C12 (ATCC), PathHunter C2C12 CXCR4 β-arrestin (DiscoverX), Ramos (ATCC), Jurkat (ATCC), K-562 (ATCC) cell lines were cultured under manufacturer recommended conditions. For flow cytometry binding assays, cells were harvested, uniquely labeled with CellTrace CSFE and Violet dyes (ThermoFisher Scientific), and pooled in equal ratios.

Clarified antibody-containing supernatant assay:

Antibody titer was first quantified using BLI. The clarified supernatant was diluted 5-fold in Octet Buffer (composed of 1X PBS, 0.1% BSA, and 0.05% TWEEN20). Octet ProteinA biosensors (Sartorius) were hydrated in supernatant from an empty vector transfection 5-fold diluted in Octet buffer and equilibrated alongside the sample plate at 30°C for 10 minutes prior to the start of the experiment. Biosensors were shaken at 1000 rpm for 60 seconds in the samples. A standard curve was created using known concentrations of VHH-Fc diluted in mammalian supernatant and Octet buffer at a 1:4 dilution. Sample binding rate over the first 60 seconds of association to the biosensor was measured. Octet Analysis Studio 13.0.2.46 software was used to build the standard curve and apply it for quantitation of samples.

0.5x10^5^ pooled cells per well were directly treated with 50 µL clarified supernatant for 2 hours at 4°C. After incubation, cells were washed four times with cold 1% PBSA, then stained with anti-Human Fc-647 secondary antibody (Biolegend) for 30 minutes at 4°C, and analyzed using a Novocyte Advanteon flow cytometer (Agilent). For PathHunter cell lines, MFI signal was compared to their respective background cell line and designs yielding MFI values greater than 3-fold over background were classified as binders.

### Purified protein assay

0.5x10^5^ pooled cells were treated with 8-point binder dilution series in 50 µL of 1% PBSA and incubated at 4°C for 2 hours. After incubation, cells were washed four times with cold 1% PBSA, then stained with anti-human Fc-647 secondary antibody (BioLegend) for 30 min at 4°C, and analyzed using a Novocyte Advanteon flow cytometer (Agilent). For PathHunter cell lines, the MFI from the respective background cell line was subtracted, and EC50s were calculated using a variable slope model in GraphPad Prism 10.

### Cell binding assay via ELISA

CXCR7 agonist candidate designs were evaluated for their ability to bind native CXCR7 on cells via cell-based ELISA. Briefly, antibody titer was quantified by BLI in clarified antibody-containing supernatants as described previously. PathHunter CHO-K1 CXCR7 β-arrestin cells (0.5x10⁵ per well) were directly treated with clarified supernatant at a final concentration of 500 nM and incubated for 2 hours at 4°C. After incubation, cells were washed four times with cold 0.1% PBSA, stained with an anti-human Fc-HRP secondary antibody (ThermoFisher Scientific) for 1 hour at 4°C, and washed again four times with cold 0.1% PBSA. SuperSignal ELISA Pico Chemiluminescent Substrate (ThermoFisher Scientific) was then added, and relative light units (RLU) was measured using an iD3 SpectraMax Microplate Reader (Molecular Devices). Designs yielding RLU values greater than 10-fold over an isotype non-binder control were classified as binders.

### Sequence novelty calculations

The sequence novelty of JAM-generated binder sequences was assessed using BLASTp searches against three databases: NR (NCBI), OAS Unpaired (OPIG), and INDI (NaturalAntibody). For each query (generated binder sequence), the top hit sequence was defined as having the highest percent identity in the query-hit alignment (pident in BLASTp output nomenclature). For each generated binder sequence, sequence novelty was defined as the percent identity between the aligned portions of the generated binder sequence and the top hit sequence.

### Structure novelty calculations

The structure novelty of JAM-generated target-binder complexes was evaluated using Foldseek queries of the target chain against the Structural Antibody Database (SAbDab). Each query returned hit complexes, where each hit complex contained a hit target chain structurally homologous to the queried target chain. The quality of each hit was assessed using combinatorial extension algorithm (CEAlign) to compute the root mean square deviation (RMSD) between the generated target chain and the respective hit target chain. Hits with target chain RMSD greater than 5 Å were excluded from further analysis.

For the remaining hit complexes, the CEAlign-derived optimal rotation matrix and translation vector were used to align the target chains of the JAM-generated complex and hit complex. The sequence similarity between the generated binder and each non-target chain in the hit complex was computed using MAFFT pairwise alignment. Hit chains that aligned with less than 50% of the residues in the generated binder sequence were excluded from further analysis. For each remaining binder-hit chain pair, the alpha carbon (C) RMSD between the MAFFT-aligned residues was calculated (without additional structural alignment, i.e., with coordinates dictated by the structural alignment of the target chains). For each generated target-binder complex, structure novelty of the binder was defined as the minimum C RMSD between the designed binder chain and a non-target chain in a hit complex (with the aforementioned constraints that the CEAlign RMSD between the target chains was at most 5 Å and at least 50% of designed binder chain aligned to corresponding hit chain).

### CXCR4 competitive inhibition assay

Binders were evaluated for their ability to block SDF1α binding to CXCR4-expressing cell lines. Briefly, Ramos and Jurkat cells were pre-treated with binders at 200nM for 2 hours at 4°C. Following incubation, 100 nM SDF1α-biotin (Protein Foundry) was added and incubated for an additional hour at 4°C. SDF1α binding was detected using a streptavidin-BV421 conjugate (BioLegend) and analyzed by flow cytometry on an Agilent Novocyte Advanteon. To quantify SDF1α blocking, MFI values were normalized to the SDF1α-only control (no binder). Percent blocking was then calculated as:

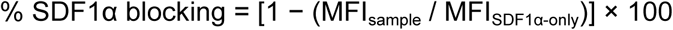

To determine antagonism potency, Ramos and Jurkat cells were pre-treated with 8-point binder titrations for 2 hours at 4°C. Following incubation, 30nM SDF1α-biotin (Protein Foundry) was added and incubated for an additional hour at 4°C. SDF1α binding was detected using a streptavidin-BV421 conjugate (BioLegend) and analyzed by flow cytometry on an Agilent Novocyte Advanteon. To quantify potency, median fluorescence intensity (MFI) values were plotted and analyzed in GraphPad Prism 10 using a variable slope model. IC50s are reported.

### CXCR7 antagonism

CXCR7 antagonism was evaluated using the PathHunter β-arrestin assay (DiscoverX), following the manufacturer’s instructions. Briefly, PathHunter CHO-K1 CXCR7 β-arrestin cells were pre-treated in triplicate with binders at 200 nM for 2 hours at 4°C. After pre-treatment, 30 nM SDF1α (Protein Foundry) was added and cells were incubated at 37°C for 90 minutes. SDF1α-induced B-arrestin signaling was detected via chemiluminescence using the PathHunter Detection Kit (DiscoverX) according to the manufacturer’s instructions.

RLU values for each sample were normalized to the RLU of the SDF1α-only control. Percent antagonism (i.e., % SDF1α signaling inhibition) was calculated as:

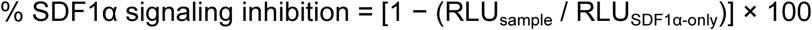

To determine antagonism potency, PathHunter CHO-K1 CXCR7 β-arrestin cells were pre-treated with 8-point binder titrations for 2 hours at 4°C. After pre-treatment, 30 nM SDF1α was added and cells were incubated at 37°C for 90 minutes. SDF1α-induced signaling was detected via chemiluminescence using the PathHunter Detection Kit according to the manufacturer’s instructions. To measure antagonism potency, RLU values for each sample were plotted and analyzed in GraphPad Prism 10 using a variable slope model. IC50s are reported with their respective 95% CI values.

### CXCR7 β-arrestin agonism

Agonism of R3 and R6 CXCR7 designs was assessed using the PathHunter β-arrestin assay (DiscoverX) according to the manufacturer’s instructions. Briefly, PathHunter CHO-K1 CXCR7 β-arrestin cells were treated with 8- or 12-point antibody titrations in duplicate for 90 min at 37°C. Following incubation, agonism signal was detected via chemiluminescence using the PathHunter Detection Kit (DiscoverX) according to the manufacturer’s instructions. RLU values for each sample were plotted and analyzed in GraphPad Prism 10 using a variable slope model. EC50s are reported with their respective 95% CI values.

CXCR7 agonist designs were screened for agonism in a single point assay. Briefly, antibody titer was quantified by BLI in clarified antibody-containing supernatants as described previously. PathHunter CHO-K1 CXCR7 β-arrestin cells were directly treated with clarified supernatant at two concentrations: the highest possible concentration and 30 nM. Cells were incubated for 90 min at 37 °C, after which agonist activity was measured as previously described. Hits were defined as designs that achieved RLU at least 3-fold greater than those of an isotype non-agonist control (08A10 VHH-Fc) at the highest tested concentration. Agonism signal at the 30 nM concentration was used to estimate the relative potency of designs by comparing their signal to that of the original agonist. 43 antibody candidates were reproduced in ExpiCHO, purified, and titrated in duplicate to evaluate agonism potency as described above.

---

**Supp. Fig. 1.**
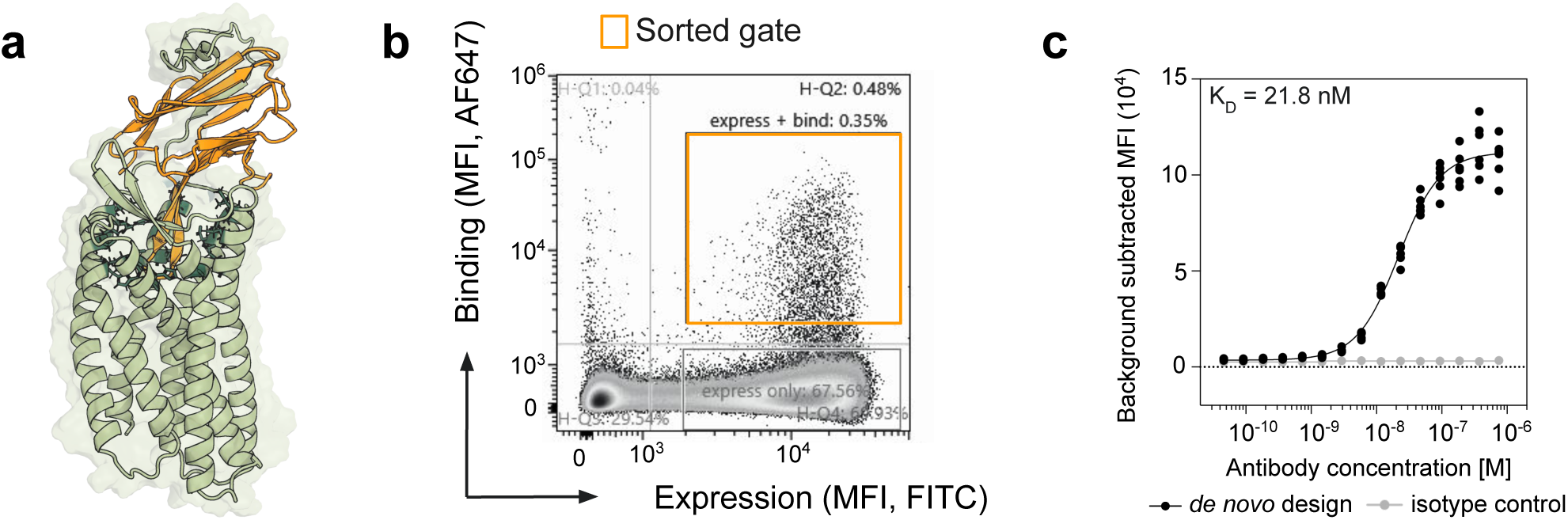
**a.** Predicted structure of *de novo* designed anti-CXCR7 antibody from R1 reported in [1] **b** FACS plots showing initial enrichments at 500 nM solCXCR7-Fc. The population within the orange gate was sorted and enriched further prior to NGS **c.** The design was expressed as an VHH-Fc in ExpiCHO supernatant (unpurified) show binding to CXCR7 overexpressed on Tango U2OS cell lines with an on-cell K_D_ of 21.8 nM.

**Supp. Fig. 2.**
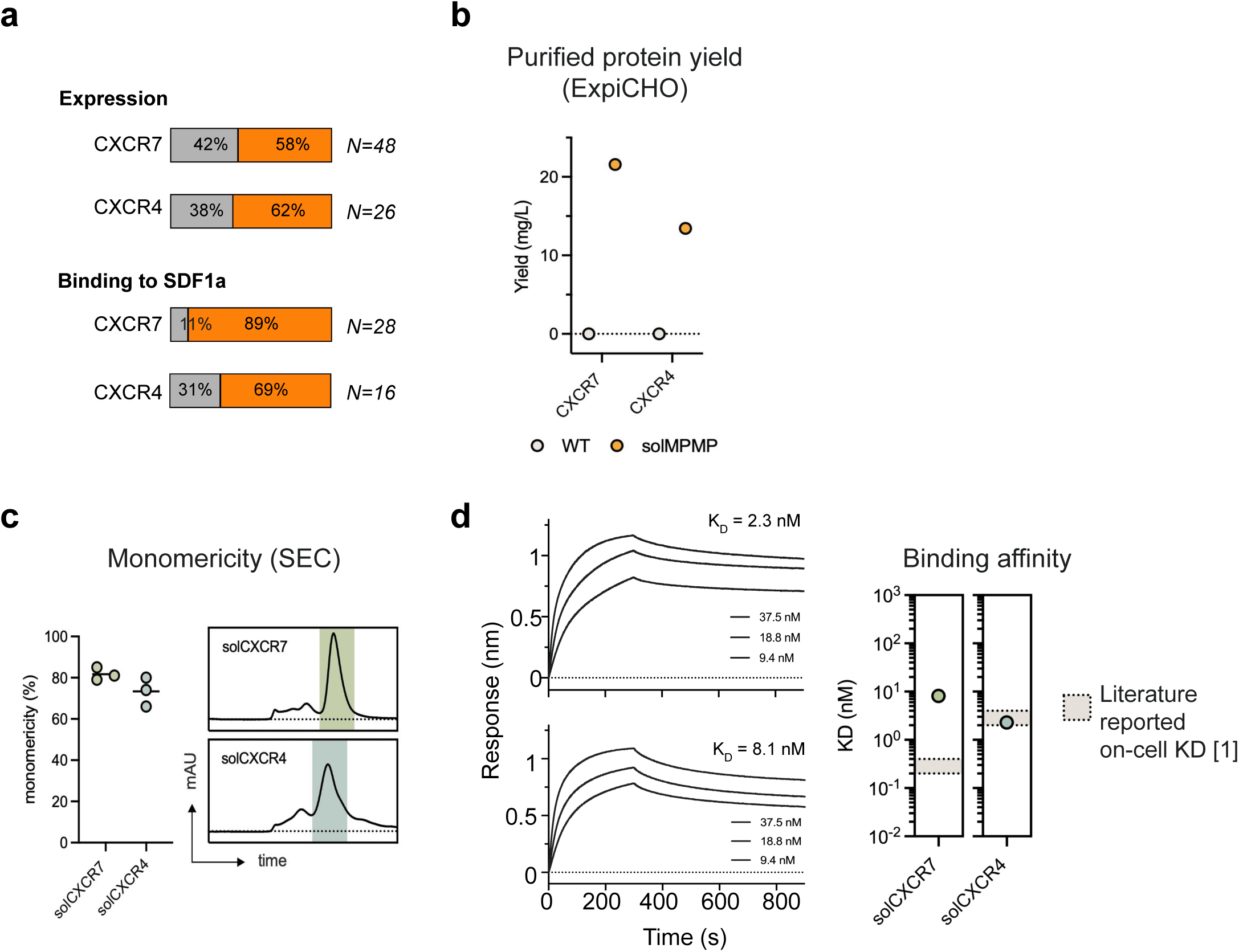
**a.** Proportion of designs that successfully expressed in ExpiCHO (top) and those that bound SDF1α in a single-point BLI (bottom). **b.** Purified protein yield of top solCXCR7 and solCXCR4 designs (orange) when produced in ExpiCHO relative to wild-type proteins (grey) **c.** solCXCR7 and solCXCR4 shows an average 82% and 73% monomericity by size exclusion chromatography (SEC) after a one-step purification; **d.** BLI-based binding studies of SDF1a to solCXCR7 and solCXCR4 show a K_D_ of 8.1 nM and 2.3 nM respectively, sufficiently proximal to reported literature values for on-cell K_D_.

**Supp. Fig. 3.**
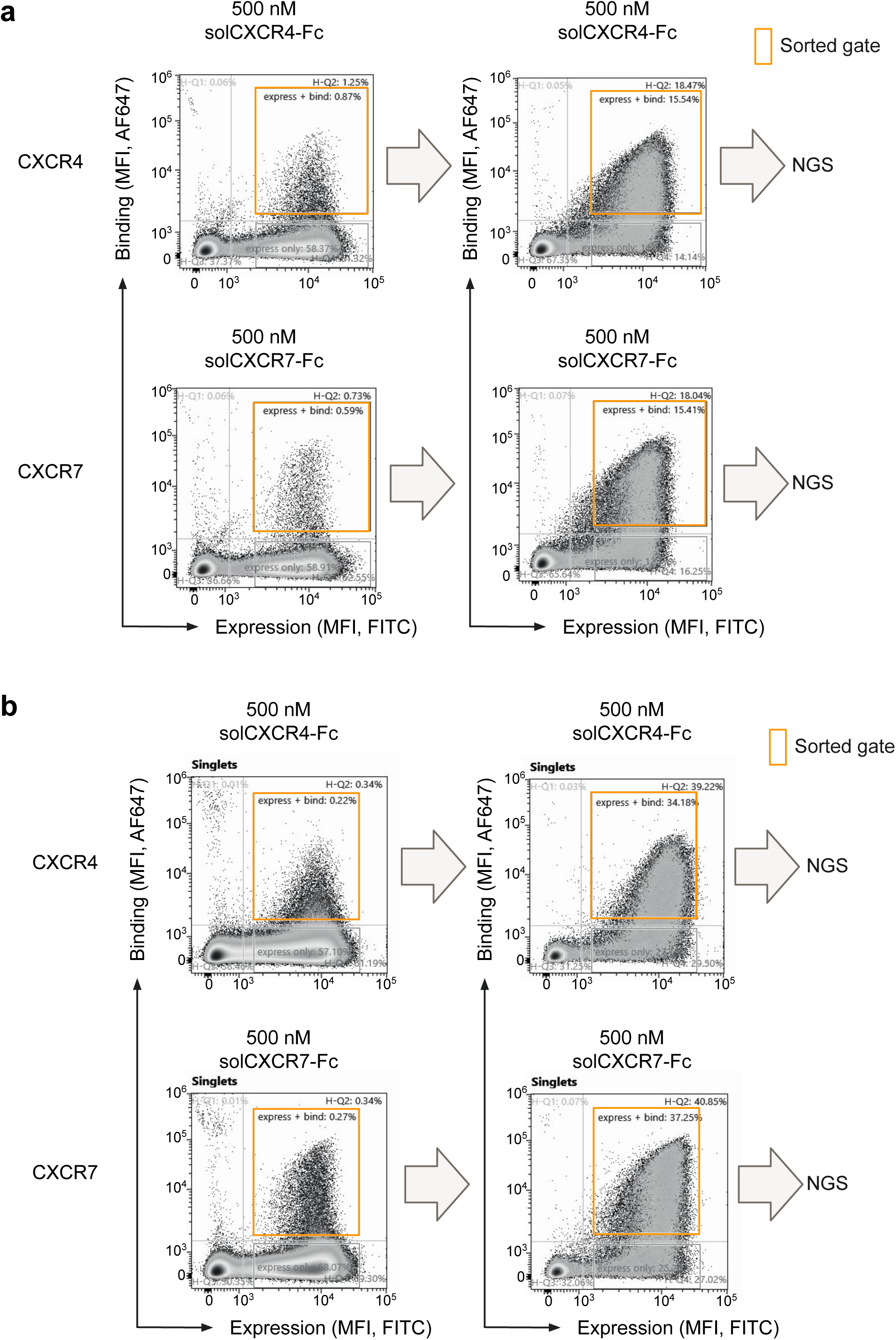
FACS plots showing initial enrichments at 500 nM for each solCXCR4 and solCXCR7 followed by a second enrichment at the same antigen concentration and NGS of double enriched populations for each (**a**) R3 and (**b**) R6.

**Supp. Fig. 4.**
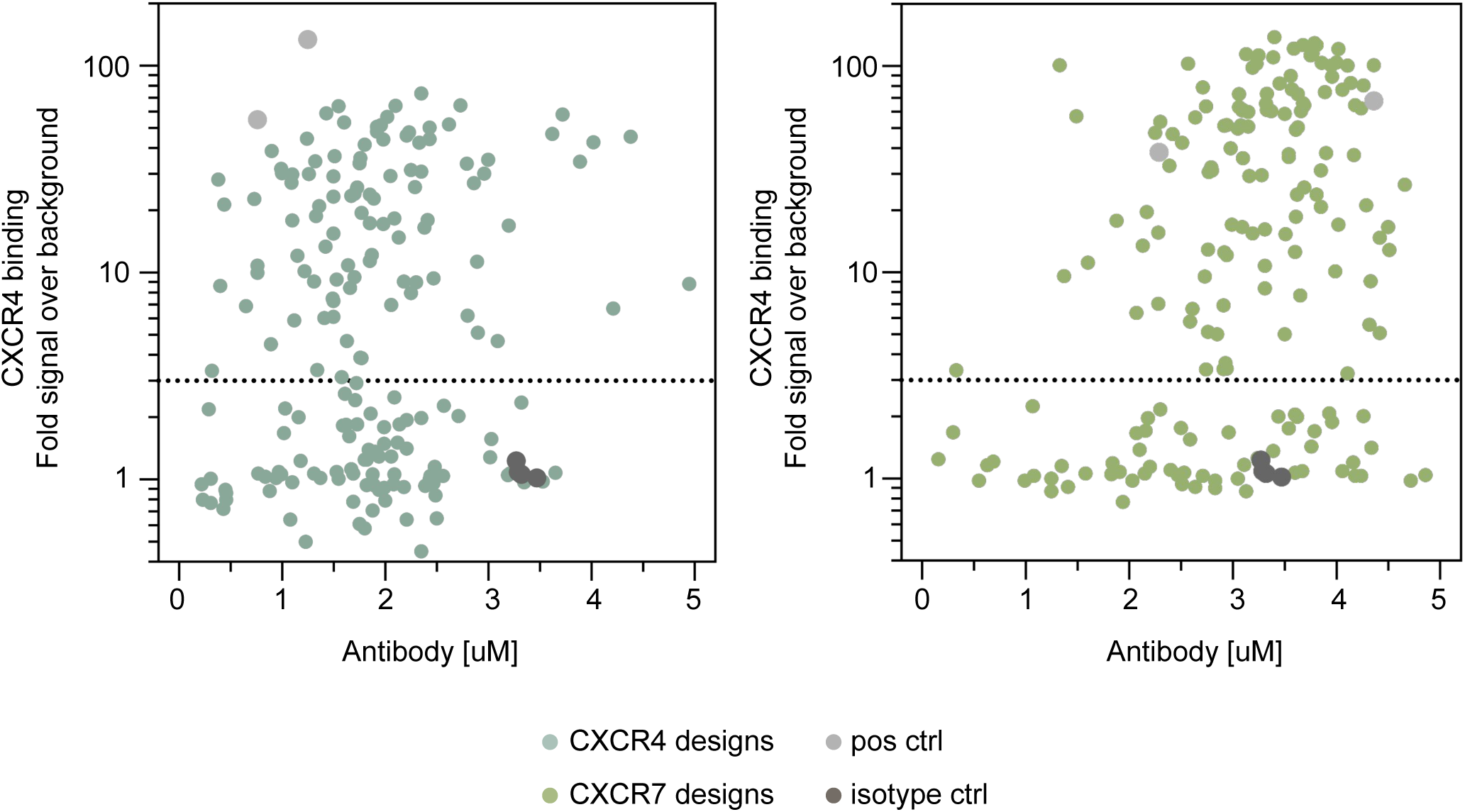
Binding determined via flow cytometry for target positive cell line over background (y-axis) vs. antibody concentration in supernatant measured by BLI (x-axis) for R6 *de novo* designs. CXCR4-targeting designs (left, blue), CXCR7-targeting designs (right, olive), and controls. Antibodies were tested at concentrations up to 5 µM. Positive controls (light gray) for each cell line and isotype controls (dark gray) are included for reference. The dotted line marks a 3-fold enrichment threshold above background.

**Supp. Fig. 5.**
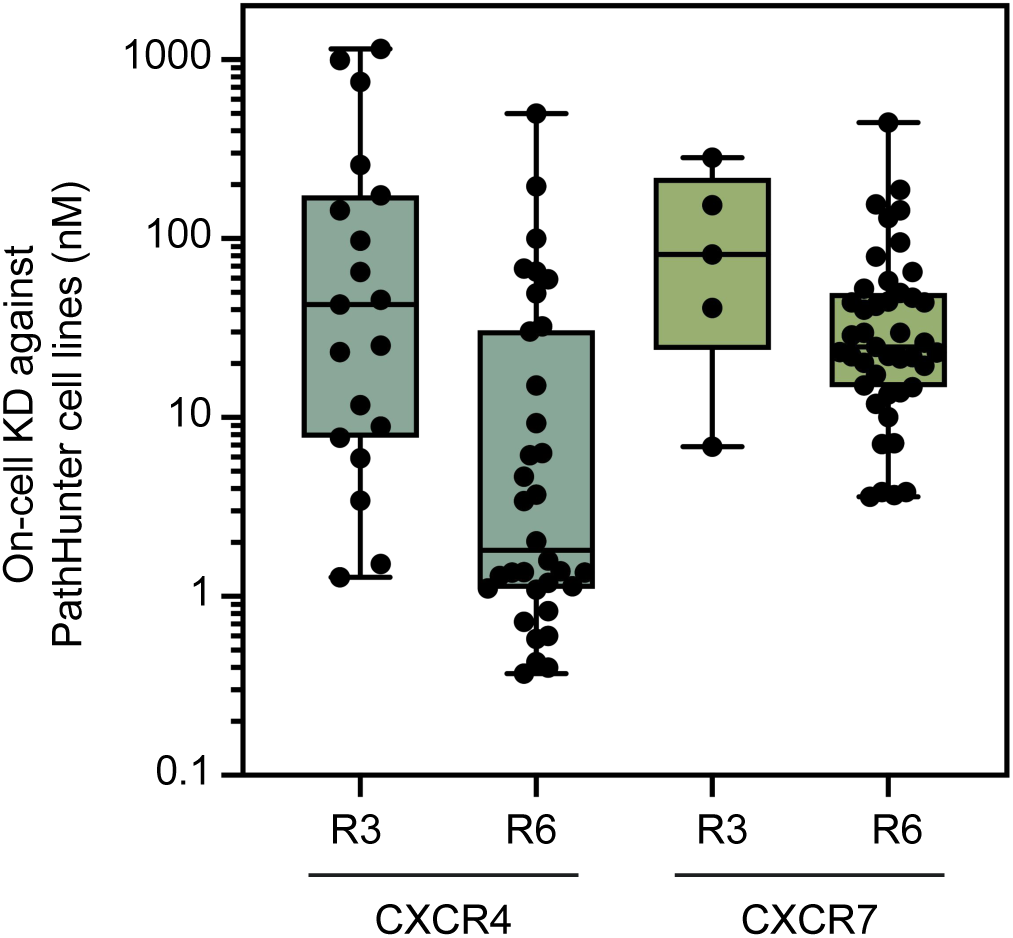
On-cell K_D_ shown for R3 and R6 designs against each CXCR7 and CXCR4. Line at median of each box plot with whiskers extending to the min. and max. Weak binders for which an accurate on-cell K_D_ was not calculable are excluded.

**Supp. Fig. 6.**
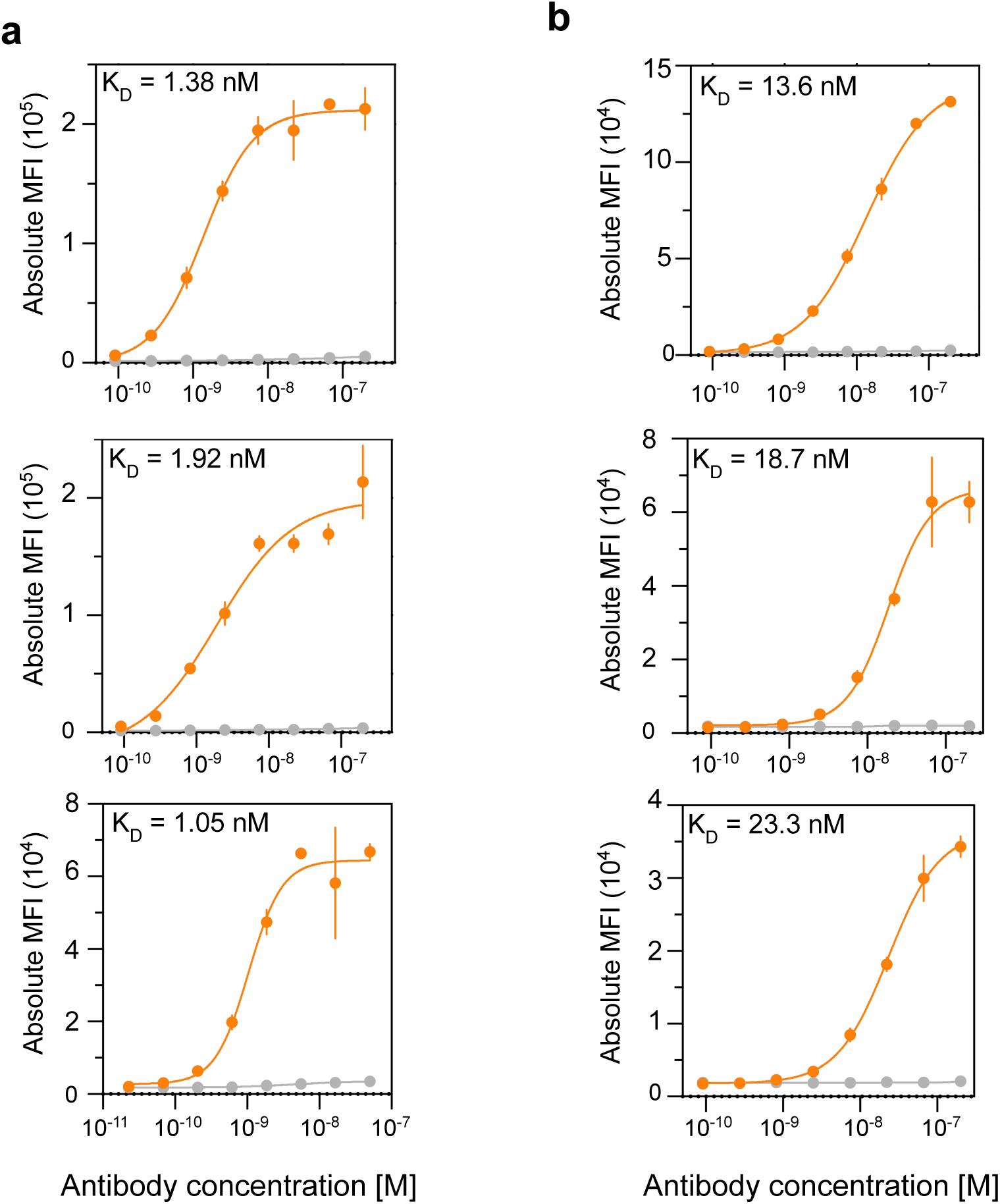
Binding trace of example **(a)** anti-CXCR4 designs and **(b)** anti-CXCR7 design against their respective PathHunter target positive (orange) and background (grey) cell line.

**Supp. Fig. 7.**
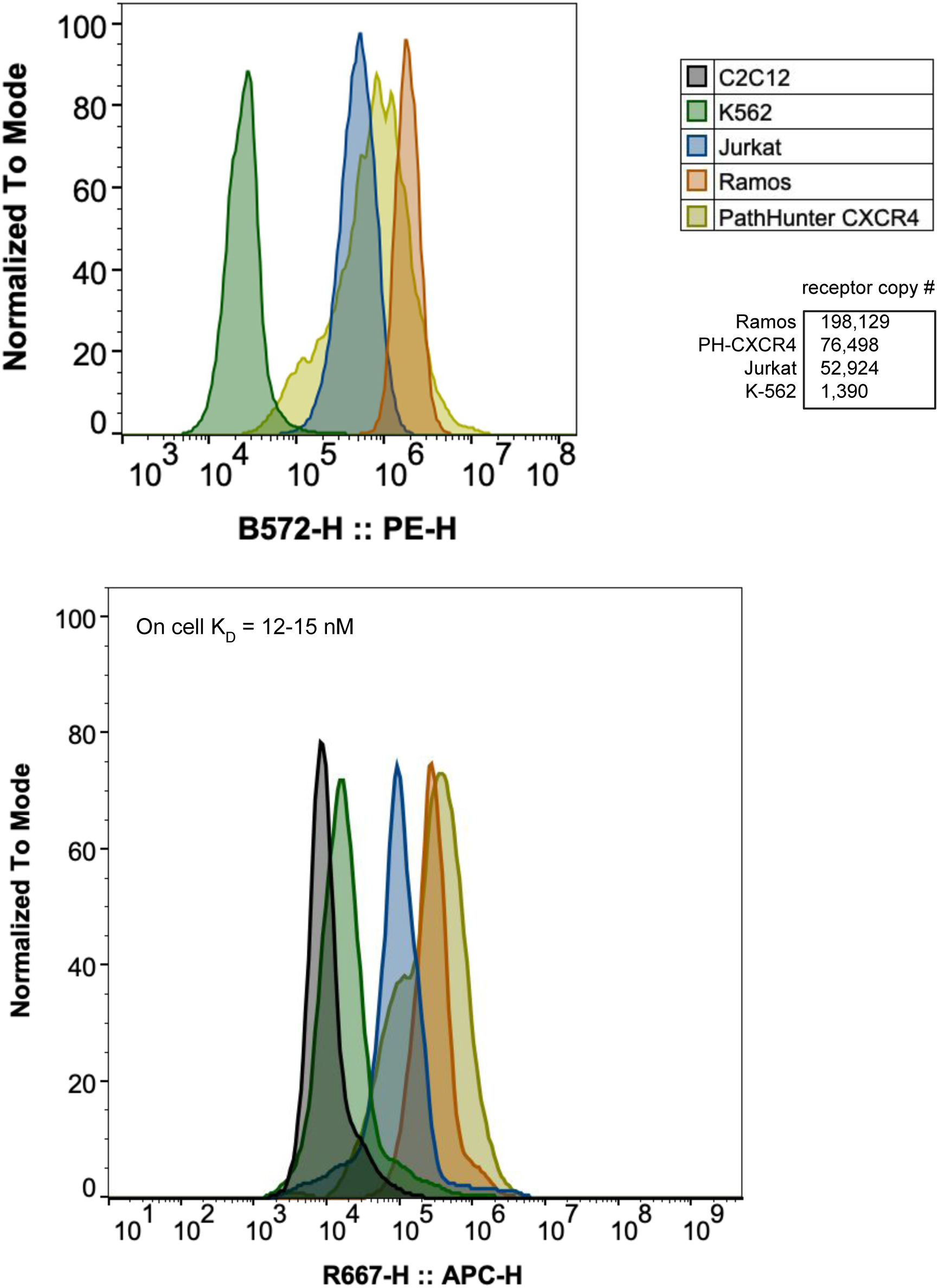
(a) PE-labeled anti-CXCR4 staining of K-562 (green), Jurkat (blue), Ramos (orange), and PathHunter CXCR4 (yellow) cells. Fluorescence is shown on a log scale and normalized to mode. Receptor copy numbers (table) were estimated using calibration beads, confirming high expression in Ramos and PH-CXCR4, moderate in Jurkat, and none in K-562 **(b)** Example binder MFI across each cell line. On-cell K_D_ for this binder ranges for 12-15 nM.

**Supp. Fig. 8.**
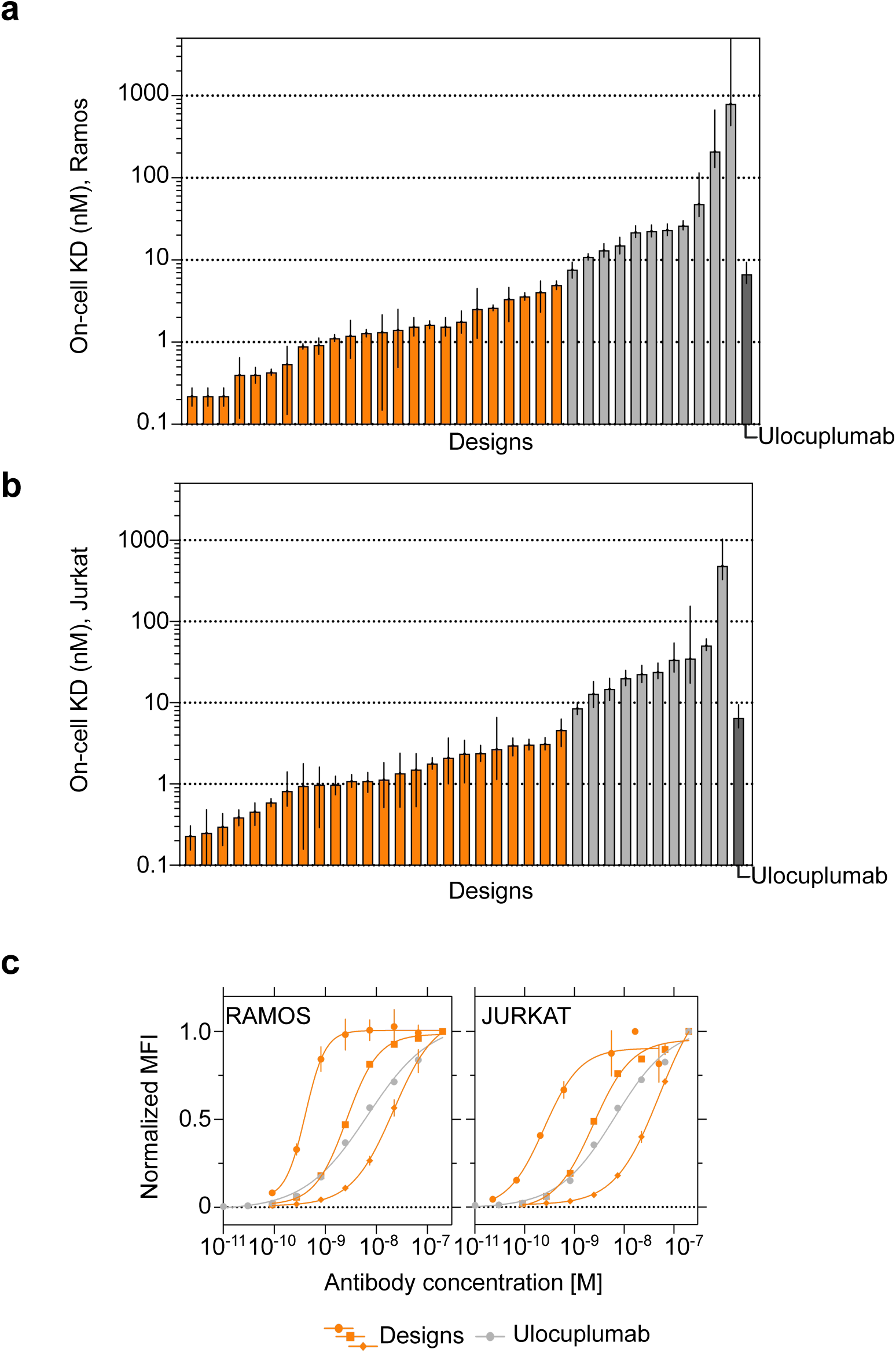
On-cell K_D_ measurements for titrations of anti-CXCR4 designs on **(a)** Ramos and **(b)** Jurkat cell lines. Error bars are 95% CI for bar plot. Orange bars denote K_D_ < Ulocuplumab. Weak binders for which an accurate on-cell K_D_ was not calculable are excluded. Binding control is for each cell line are shown in dark grey. **c.** Binding trace of best anti-CXCR4 designs on Ramos (left) and Jurkat (right). Error bars are SD.

**Supp. Fig. 9.**
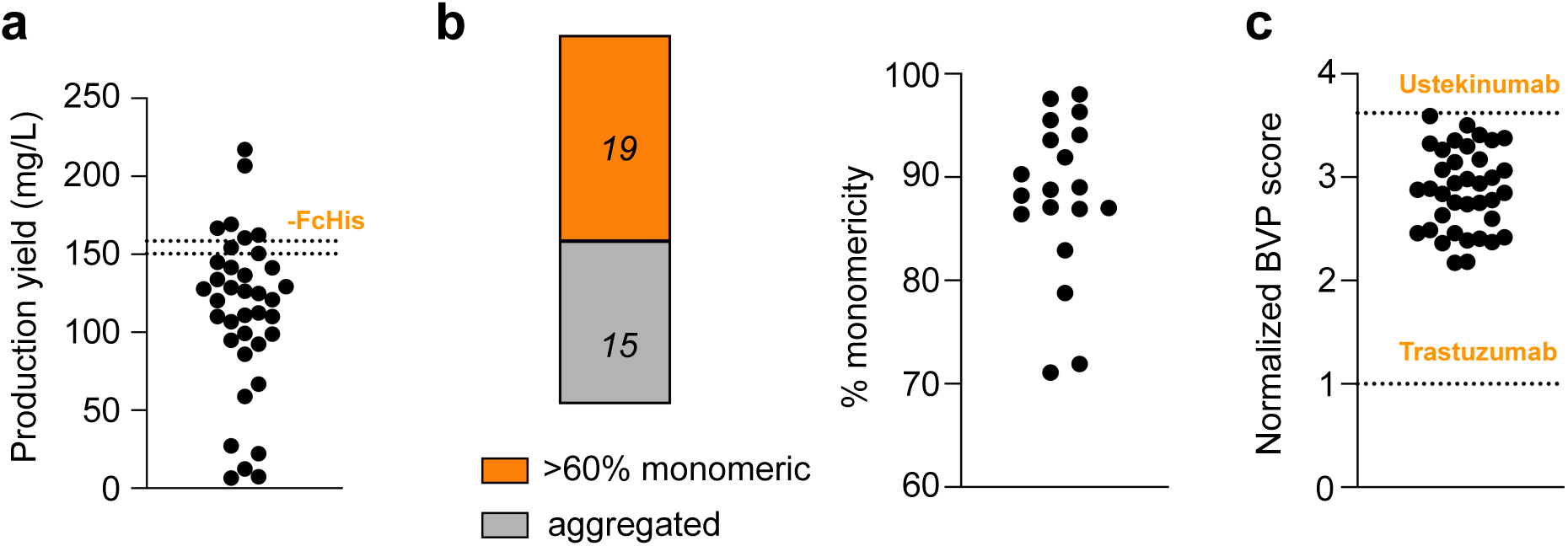
Early stage developability of anti-CXCR4 R6 designs. **a.** Production yield from 24-well ExpiCHO production where the dotted lines indicates production yield for Fc-His produced concurrently in duplicate; **b.** SEC shows binder monomericity upon a one-step high-throughput Protein A-based purification. Binders with monomercity <60% are classified as aggregated; **c.** Polyspecificity scores by BVP ELISA and normalized to Trastuzumab (lower orange line at y=1; upper orange line is Ustekinumab assayed concurrently)

**Supp. Fig. 10.**
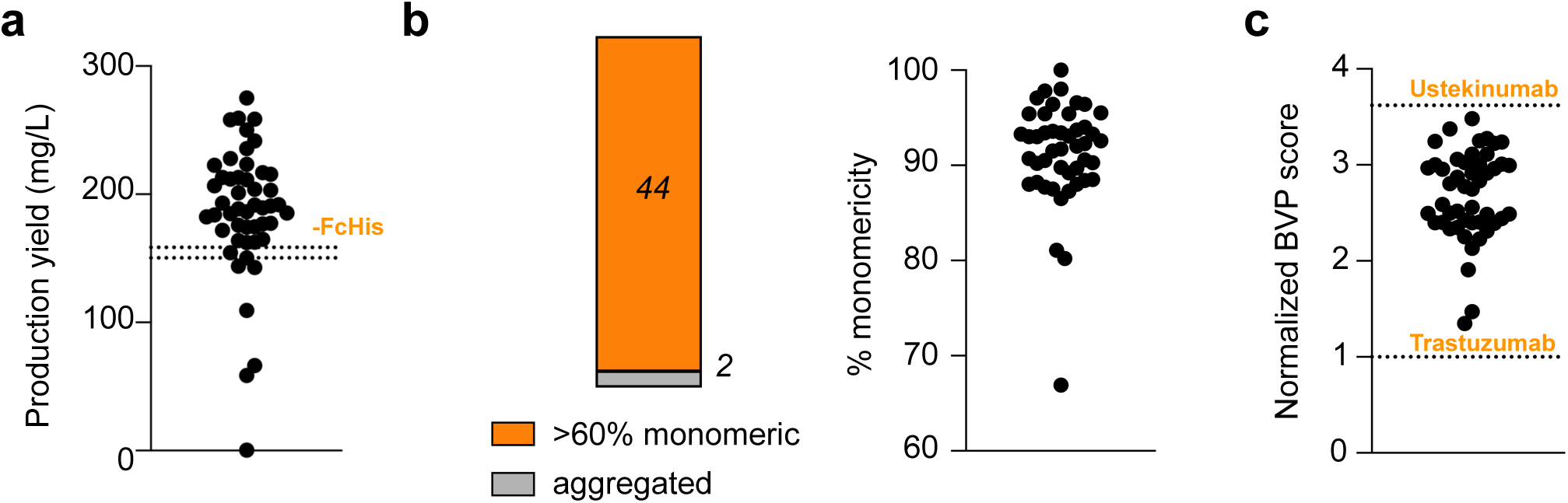
Early stage developability of anti-CXCR7 R6 designs. **a.** Production yield from 24-well ExpiCHO production where the dotted line indicates production yield for -FcHis produced concurrently in duplicate; **b.** SEC shows binders monomericity upon one-step high-throughput Protein A-based purification. Binders with monomercity <60% are classified as aggregated; **c.** Polyspecificity scores by BVP ELISA and normalized to Trastuzumab (lower dotted line at y=1; upper dotted line is Ustekinumab assayed concurrently)

**Supp. Fig. 11.**
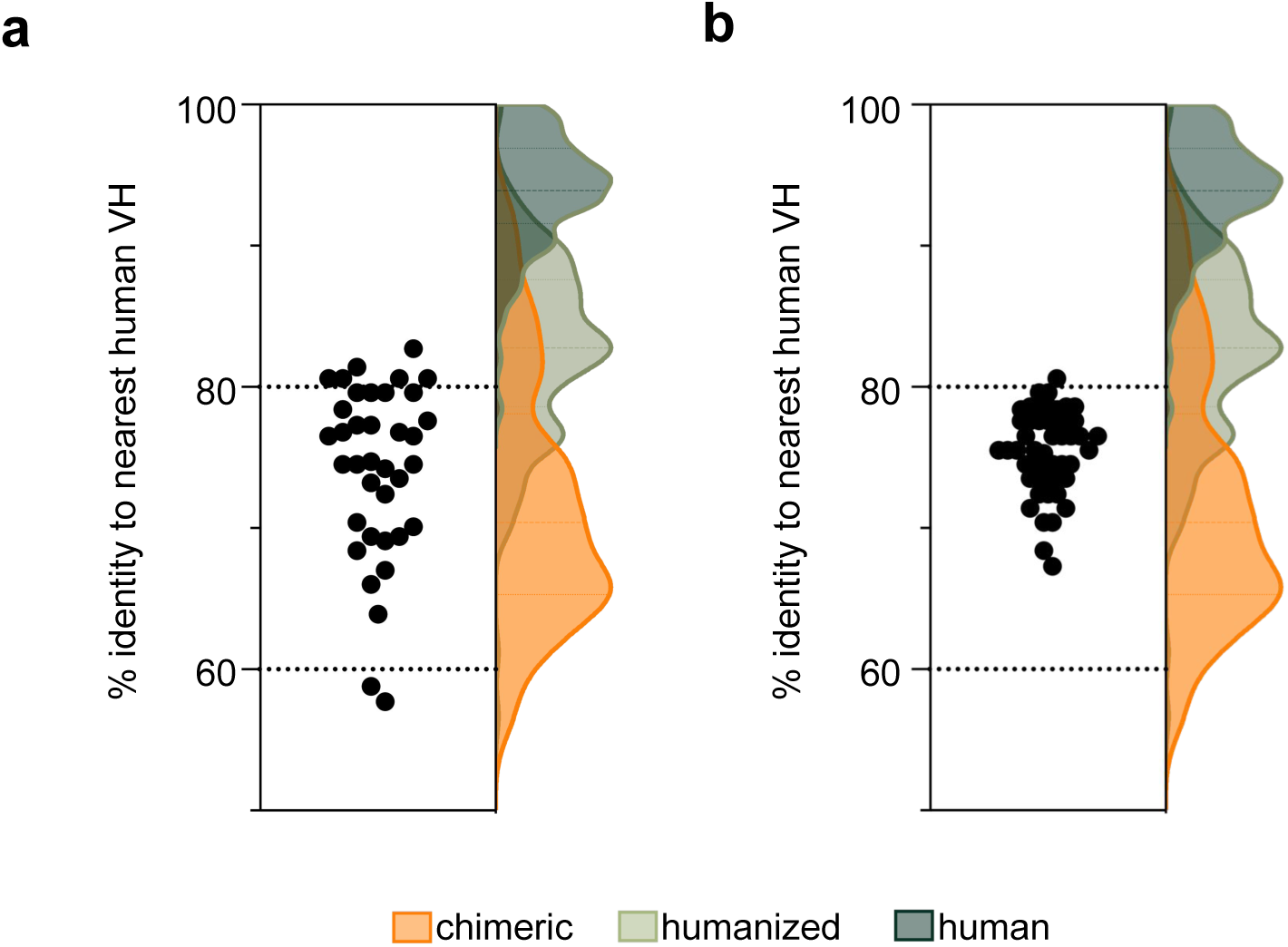
Percentage amino acid sequence identity to nearest human germline for VH sequences for **(a)** anti-CXCR4 R6 designs and **(b)** anti-CXCR7 R6 designs. Colored kernel density estimates outside of the stripplot indicate % identity to nearest human germline values for clinical stage chimeric, humanized, and human antibodies deposited in Thera-SAbDab [1].

**Supp. Fig. 12.**
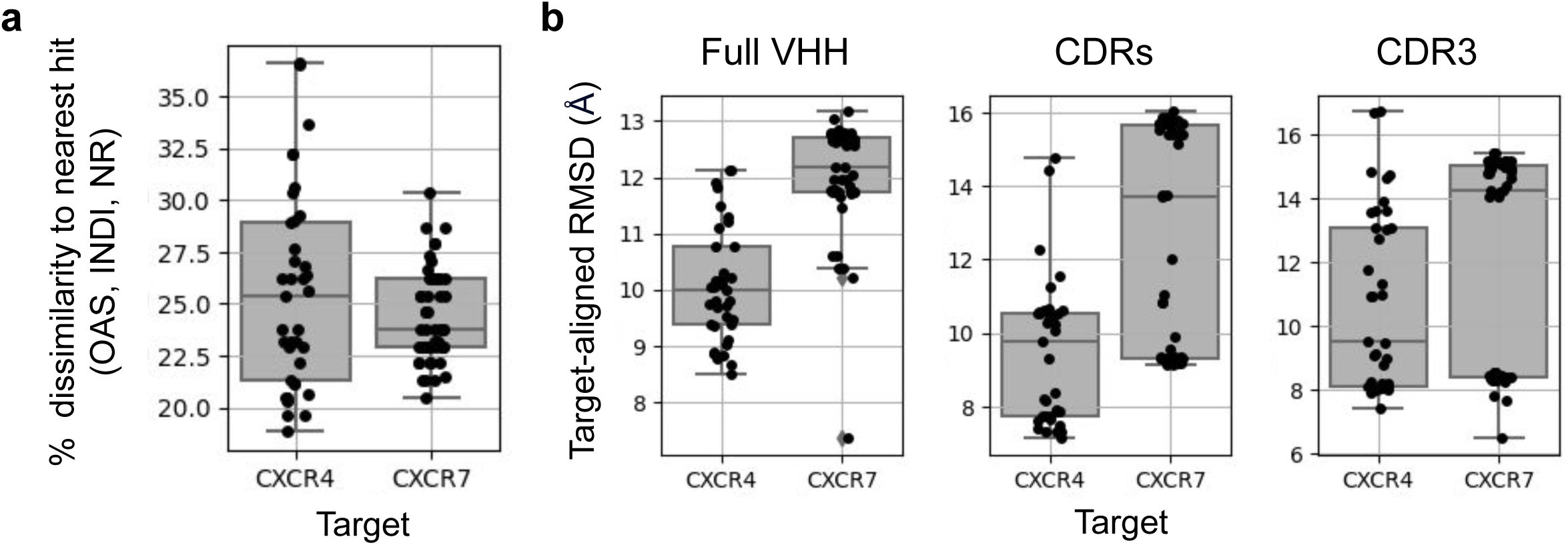
Sequence and structure novelty. **a.** Sequence novelty as measured by percent identity to nearest BLAST hit from a combined sequence database containing all sequences from the OAS (OPIG), INDI (NaturalAntibody), and NR (NCBI) databases (>3 billion sequences total). **b.** Structure novelty for each *de novo* VHH-GPCR complex as measured by alpha-carbon RMSD of the full VHH structure (left), all CDRs (middle), CDR3 region (right) to the binder in the most similar binder-target complex structure in SAbDab (OPIG). Briefly, the target chain of the JAM-generated VHH-target complex was used as a query to a FoldSeek-based search of SabDab. For each hit, the JAM-generated VHH-target complex and the hit complex were aligned on the target chain and the RMSD between the JAM-generated VHH and binder chain the hit complex was calculated. The minimum RMSD structure across hits was taken as the most similar binder-target complex structure. See Methods for full implementation details on sequence and structure novelty calculations.

**Supp. Fig. 13.**
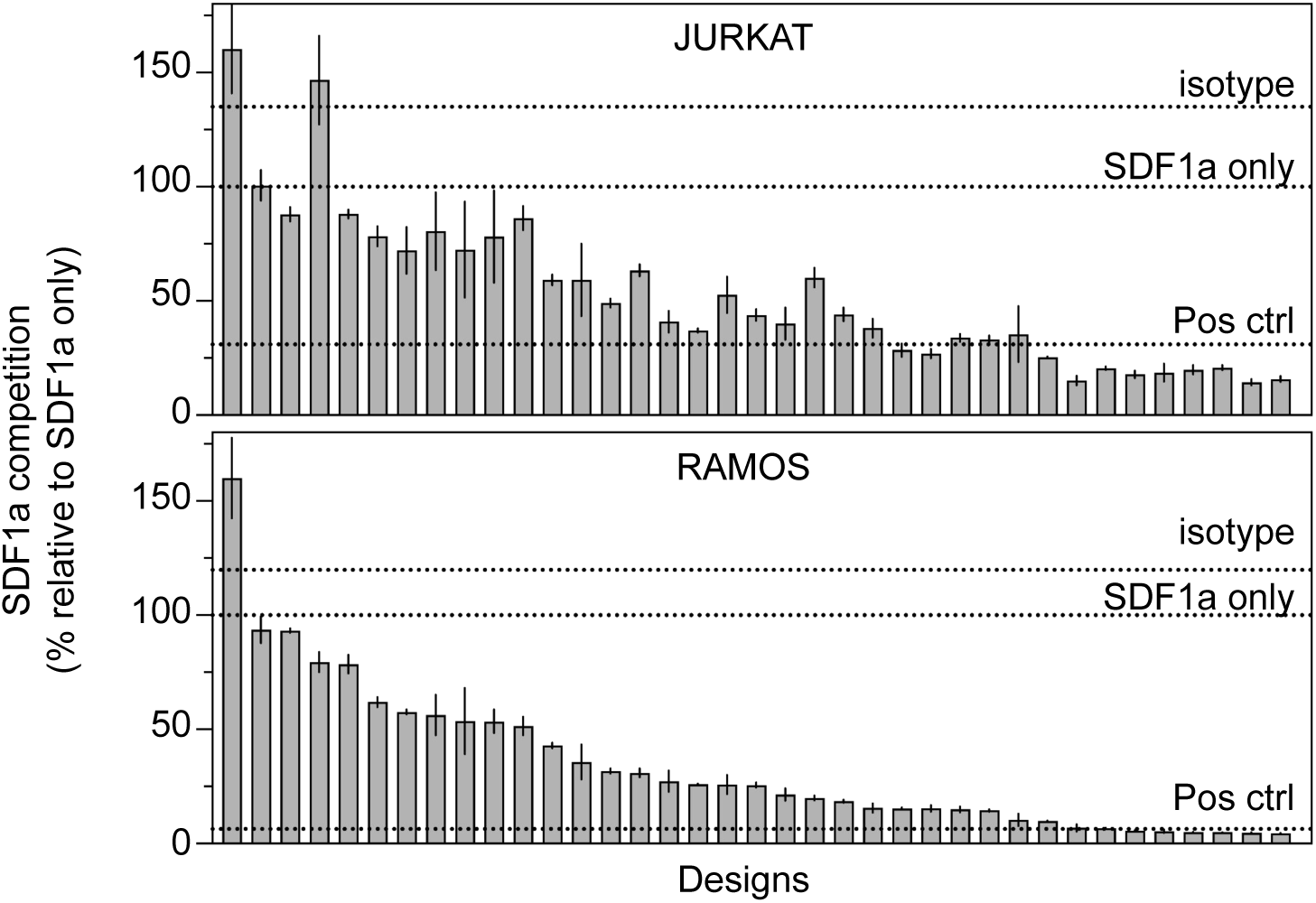
Jurkat (top) and Ramos (bottom) cells were pre-treated with 200 nM of each purified antibody binder for 2 hours at 4°C, followed by incubation with 100 nM SDF1α. SDF1α binding was measured by flow cytometry, and competition was calculated as percent binding relative to the SDF1α-only condition. Isotype and positive control antibody 238D2, a known antagonist, are included as controls [1]. Error bars represent standard deviation across triplicates.

**Supp. Fig. 14.**
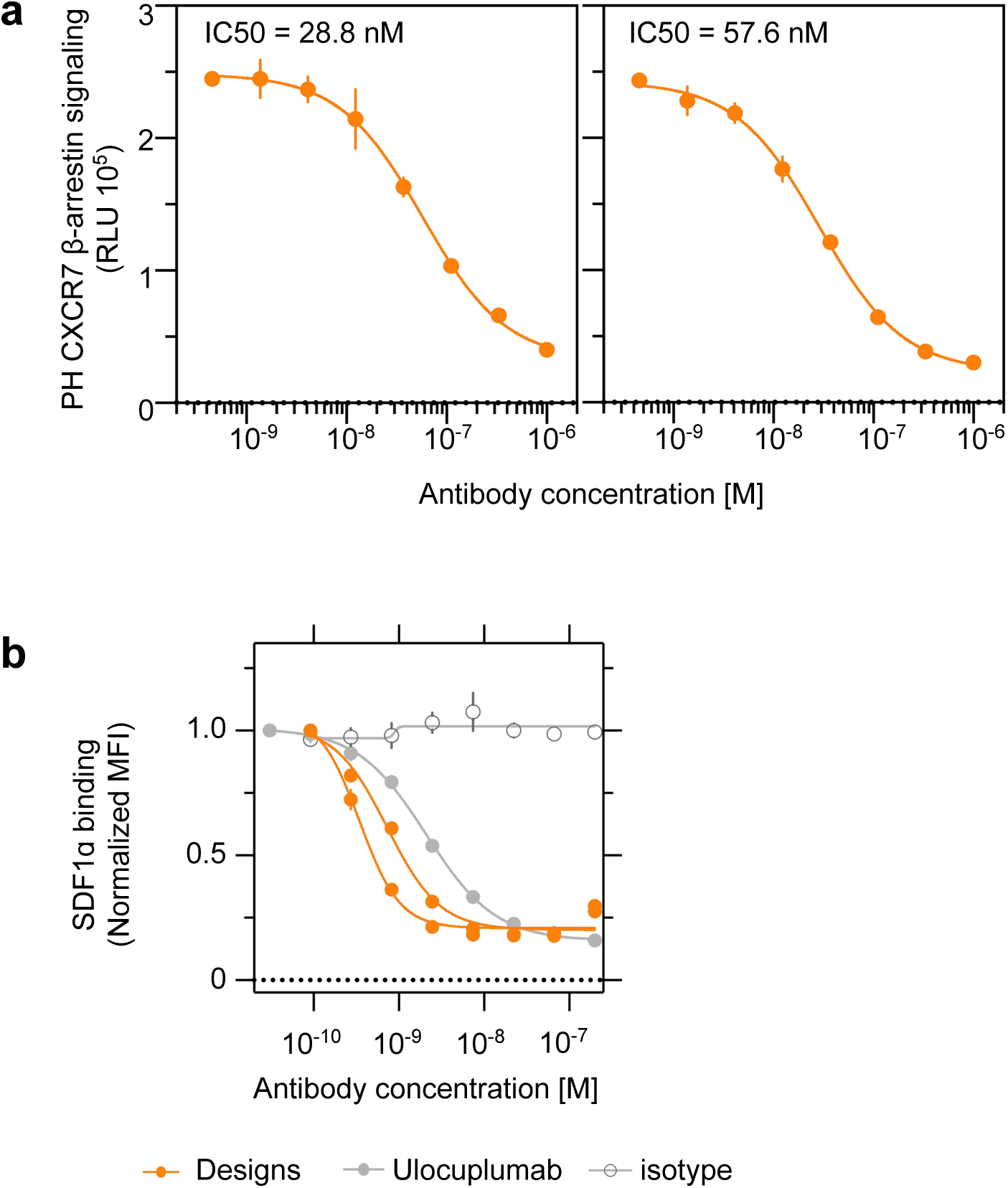
**a.** Binding trace of antagonistic potency (IC50) of two anti-CXCR7 designs against SDF1α-induced β-arrestin signaling. Error bars are SD. **b.** Flow cytometry-based titrations show displacement of fluorescently labeled SDF1α from Jurkat cells by designs (orange), Ulocuplumab (light grey) and an isotype control (grey, hollow symbol).

**Supp. Fig. 15.**
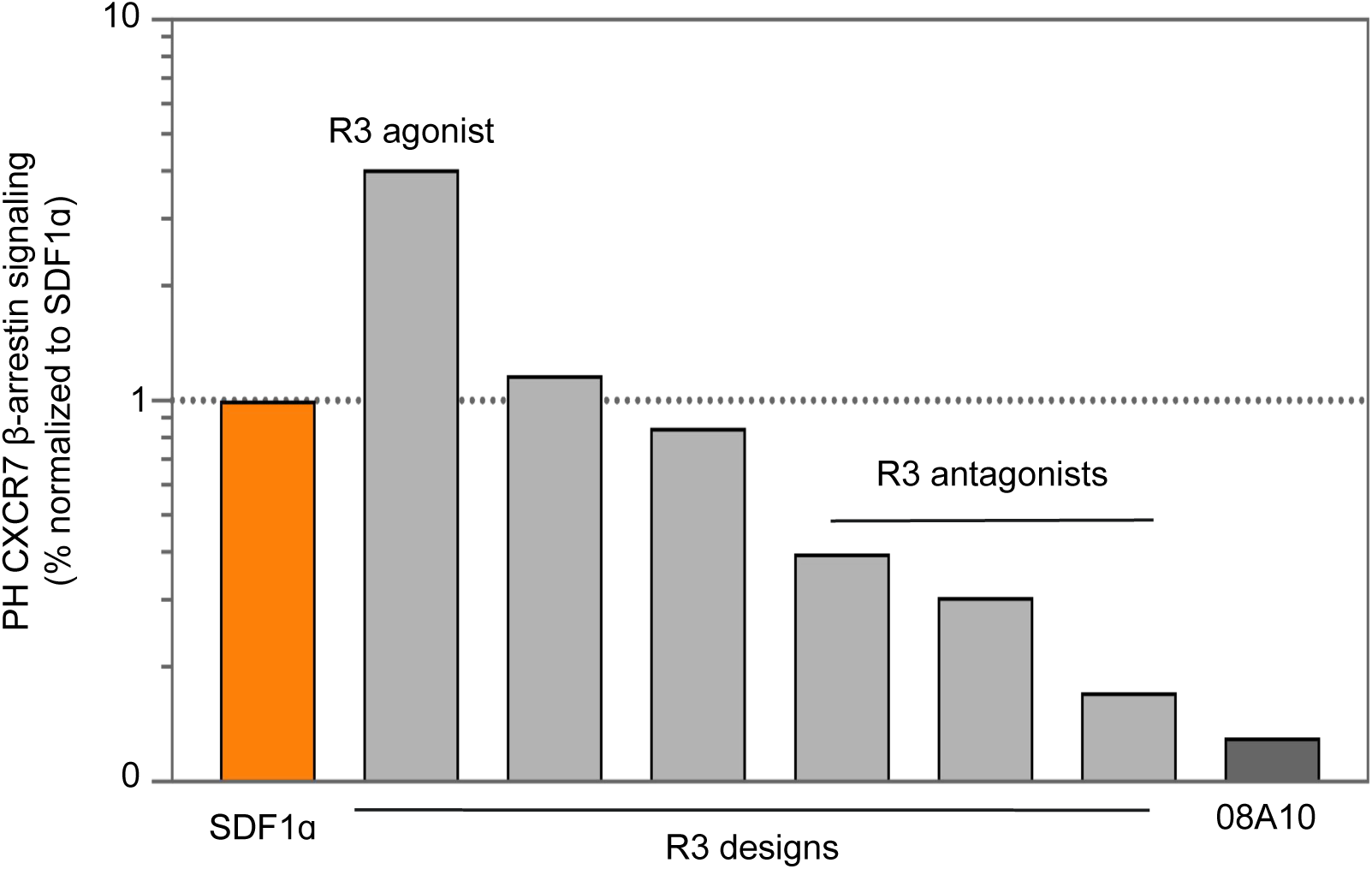
β-arrestin signaling on PathHunter CXCR7 cells in response to R3 antibody designs in conjunction with SDF1α, normalized to SDF1α-only. Designs span both agonists and antagonists. SDF1ɑ (orange) and pos ctrl (08A10, dark grey) were assayed concurrently. incubated overnight at 37°C. The following day, cells were pre-treated with 5 µL of antagonist (10X in PBS; final assay concentration = 1 µM) for 30 minutes at 37°C. Subsequently, 5 µL of SDF1α agonist (10X EC80 in PBS; final assay concentration = 7 nM) was added to each well and cells were incubated for 90 minutes at 37°C. Agonist EC₈₀ values were pre-determined for each receptor (CXCR7: 7 nM). Finally, 50 µL of working detection solution was added, bringing the total assay volume to 150 µL. Luminescence was measured per the manufacturer’s protocol. All conditions were run in triplicate.

